# Engineering *in vitro* models of skeletal muscle with neuromuscular junctions using hierarchical micro-nano biomaterials: Cooperative effect of adhesion ligand nanoclustering and surface anisotropy

**DOI:** 10.64898/2026.04.13.717344

**Authors:** Shirin Nour, Kristy Swiderski, Sahar Salehi, Gordon S. Lynch, Andrea J. O’Connor, Greg G. Qiao, Daniel E. Heath

**Author notes:** Corresponding author: Associate Professor Daniel Heath.

## Abstract

Robust development of *in vitro* mature skeletal muscle with functional neuromuscular junctions is an unmet challenge that must be addressed for advances in skeletal muscle tissue engineering and for the development of skeletal muscle tissue models for disease modelling and drug discovery. Herein, we developed hierarchical, anisotropic biomaterials that induced early maturation of more mature myotubes and the development of neuromuscular junctions (NMJs) during co-culture with motor neurons. We accomplished this by creating micro-nano biomaterial interfaces that presented nanoclusters of integrin-binding ligands to promote mechanotransduction on the surface of aligned electrospun microfibers. Controlling surface topography and nanoscale ligand clustering led to 1.5 to 2.5-fold increases in myoblast proliferation, myotube formation, elongation, and alignment; resulting in spontaneous twitching; and enhanced myotube-neuron connections, including increasing acetylcholine receptor clustering, neurite branching, and myotube contraction compared to control surfaces. These findings highlight the importance of tailoring adhesive peptide distribution and presentation for the *in vitro* development of NMJs and synaptic organization. This approach offers a valuable platform for fundamental research on muscle development and neuromuscular diseases, paving the way for improved skeletal muscle tissue engineering and drug screening strategies.

## 1. Introduction

Skeletal muscle is a hierarchical and structurally ordered tissue made from aligned muscle fiber bundles surrounded by vasculature and neurons ^[1,2]^. Loss of skeletal muscle function due to musculoskeletal disorders is the second leading cause of disability globally and is responsible for more than $380 billion per annum in healthcare costs in the United States alone ^[3]^. Tissue engineering strategies have the potential to restore skeletal muscle function and to create skeletal muscle-on-a-chip devices for disease modeling and drug discovery ^[4,5]^. However, existing technologies are unable to rapidly create directionally aligned and mature myotubes that are innervated with motor neurons, limiting progress in the field ^[6,7]^.

The neuromuscular junction (NMJ) is a specialized synapse between motor neurons and skeletal muscle fibers, facilitating muscle contraction and motor functions, such as movement, respiration, and speech ^[8]^. Degenerative muscle diseases with dysfunctional NMJs can compromise muscle mass, organization, and elasticity, leading to muscle atrophy and loss of contractility ^[9,10]^. Limitations of current treatments for skeletal muscle injuries and neuromuscular diseases necessitate deeper knowledge about the biochemical mechanisms underlying degeneration of cellular interconnections and the development of biomaterial-assisted *in vitro* models, particularly for studying NMJ pathophysiology and for drug screening^[11,12]^.

Previous studies from our group and others highlighted the crucial role of integrin-mediated mechanotransduction in activation of Ras homolog family member A (RhoA) and focal adhesion kinase (FAK) signaling pathways, which regulate myoblast proliferation, migration and alignment during myogenesis, facilitating fusion into multinucleated myotubes and proper function and integrity during contraction, and we have developed biointerfaces that promote focal adhesion formation through nano-scale clustering of integrin-binding ligands ^[13–17]^.

Additionally, integrins are involved in the differentiation, assembly, and maintenance of the structural integrity of the postsynaptic membrane at the NMJ ^[2,18,19]^. Integrins in muscle fibers and presynaptic membrane are involved in organizing the actin cytoskeleton, which contributes to the clustering of acetylcholine receptors (AChRs) at the synaptic cleft and the localization of active zone components with efficient synaptic vesicle release, facilitating efficient neurotransmission for muscle contraction ^[16,20]^.

In addition to clustering and efficient engagement of cell adhesion ligands on biomaterial surfaces, providing physical cues such as surface anisotropy (e.g., in the form of aligned fibers or microchannels) significantly influences myoblast behavior ^[21]^. These cues can direct the orientation and elongation of myoblasts into aligned structures and promote fusion into mature myotubes, better resembling the native architecture of muscle tissue ^[22,23]^. Furthermore, changes in the arrangement of the cytoskeleton in response to surface topography may positively impact integrin-mediated mechanotransduction, receptor clustering, and the strength of developed focal adhesion ^[23]^.

Engineering a biointerface’s biochemical and physical properties to control adhesion molecules’ density and patterning, and substrate topography, could be a promising tool in generating biomimetic skeletal muscle tissue models ^[24,25]^. Despite the growing interest in developing NMJs *in vitro*, this is the first study utilizing advanced anisotropic biomaterials with controlled nanoscale presentation and distribution of integrin binding peptides for skeletal muscle tissue regeneration and NMJ formation, to our knowledge. Additionally, this biomaterials-based strategy has significant advantages over competing technologies because it does not require expensive and complex equipment/manufacturing techniques, such as electrical stimulation bioreactors ^[26–28]^.

In this study, we hypothesized that integrating microarchitectural anisotropy and nanoclustering of cell adhesion cues would create a supportive environment for myogenesis and de novo formation of NMJs. Our approach using murine myoblasts co-cultured with mouse motor neurons on electrospun RGD-functionalized random copolymers, explored peptide nanocluster density and microfiber alignment on myoblast behaviors, and myogenesis, followed by neuromuscular connection. In addition to enhancing myogenic differentiation, neurite length, and NMJ formation and function, the results highlight the potential of our system for modelling skeletal muscle tissue regeneration for drug screening related to neuromuscular degeneration.

## 2. Results and discussion

### 2.1. Synthesis, fabrication, and characterization of hierarchical biointerfaces with nano-scale clustering of cell adhesive ligand and anisotropic micropatterning

The primary goal of this research is to improve the *in vitro* development of engineered skeletal muscle and enhance its functional connection to motor neurons through the development of neuromuscular junctions. To accomplish this, we synthesized a family of random comb co-polymers that can be fabricated into planar and micropatterned surfaces with controlled presentation of cell adhesive ligands ^[15,29]^. The polymers were characterized by ^1^H NMR and GPC analysis. The first polymer is a co-polymer of methyl methacrylate (MMA) and poly(ethylene glycol) methacrylate (PEGMA, M_W_ = 475 g.mol^-1^) with a number average molecular weight of 128 kDa and a polydispersity index of 1.1, hereafter referred to as the MP polymer (**Figure S1** and **Table S1**). We also produced a terpolymer from methyl methacrylate (MMA), poly(ethylene glycol) methacrylate (PEGMA, M_W_ = 475 g.mol^-1^), and a norbornene-functionalised poly(ethylene glycol) methacrylate (PEGMA, M_W_ = 668 g.mol^-1^) with a number average molecular weight of 182 kDa and a polydispersity index of 1.2, hereafter referred to as MPP polymer (**Figure S2**). The norbornene groups were subsequently reacted with cysteine-modified RGD ligands to incorporate integrin-binding properties to the material according to our previously published protocol (**Figures S2**, **Figure S3**, and **Table S1)**^[15]^.

Bulk materials made from the comb polymers are water stable due to the hydrophobic MMA repeat units, yet their surfaces possess robust non-fouling properties due to the hydrophilic PEG pendant groups, making them an ideal background to study how ligand functionalization impacts cellular processes ^[15,29–31]^. Additionally, the materials are soluble in a variety of organic solvents, enabling facile fabrication of complex geometries through solution processing. We capitalized on these properties to make a library of biointerfaces with control over the ligand presentation and micropatterning, as demonstrated in **Figure 1**.

**Figure 1.**
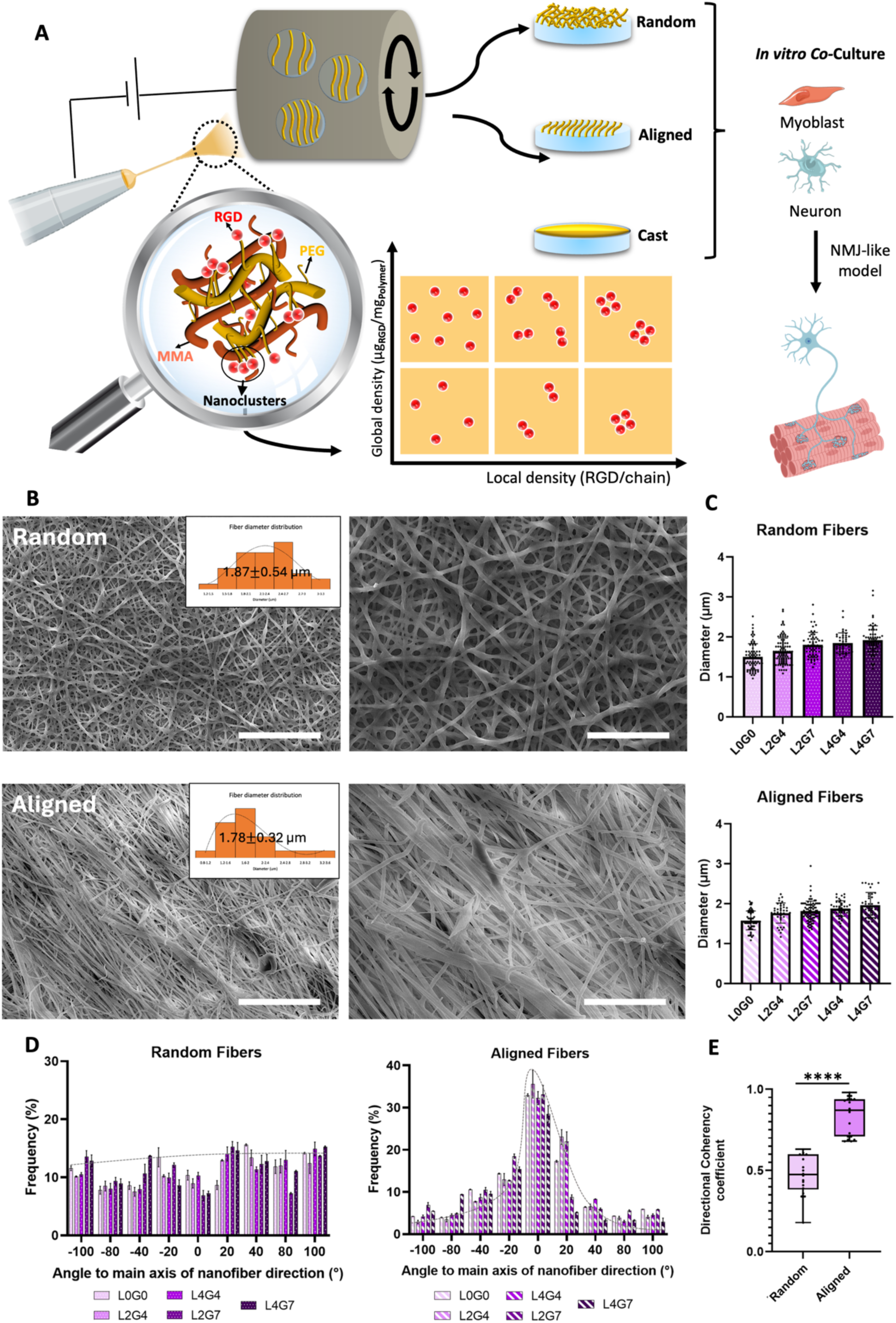
Schematic demonstrating fabrication of micropatterned biointerfaces with controlled presentation of the cell-adhesive RGD ligands for use in skeletal muscle and NMJ formation *in vitro*. A) MP (orange) and RGD-functionalized MPP (yellow) polymers were blended and fabricated into cast films or random or aligned microfiber biointerfaces with controlled local and global ligand densities. B) Scanning electron microscopy (SEM) confirms the random and aligned fiber architectures, and the histograms show the distribution of fiber diameters. Scale bars are 100 and 50 µm for low and high-magnification SEM images, respectively. C) Quantification of average fiber diameter demonstrates that it was constant across all formulations, with no statistical differences between samples (n = 100 measurements). D) Quantification of fiber alignment and E) directional coherency coefficient demonstrate a significant increase in orientation of the fibers in the direction of rotation in aligned samples compared to random fibers (n = 10). Asterisks (*) above the bars indicate statistical differences between groups, **** represents a p-value of ≤ 0.0001.

Experimental and modelling work has demonstrated that in bulk materials, the comb polymers at the interface adopt a quasi-2D conformation, where the hydrophobic polymer backbone lies in the plane of the surface and the hydrophilic PEG pendant groups partition to the aqueous phase ^[29,32]^. This unique property allows the global ligand density (the total amount of ligand in the material) to be controlled by controlling the blending ratio between MP and MPP polymers, and it allows for the local density (the degree of nanoscale clustering of ligands) to be controlled by controlling the degree of functionalisation of the MPP chains. In this work, we created polymer blends with a range of local RGD ligand densities (L) from 0-4 peptides per polymer chain and a range of global ligand densities (G) from 0-7 µg_peptide_/mg_polymer_, as compiled in **Table 1**. Biointerfaces made from the unfunctionalized MP polymer and those functionalised with the non-adhesive RGE peptide were used as controls.

**Table 1.** Polymer samples used in this study to generate cell culture surfaces.

| Sample name | Polymer | Local ligand density (ligands/MPP polymer chain) | Bulk ligand density of MPP polymer ($\mu\text{g peptide}/\text{mg polymer}$) | Global ligand density of MP+MPP polymer blend ($\mu\text{g peptide}/\text{mg polymer}$) | Molar blending ratio (MPP/MP) | PEG content (mol%) |
| --- | --- | --- | --- | --- | --- | --- |
| **L0G0** | MP | - | - | - | - | 15.6 |
| **L1G7** | MPP | 1 | 7.5 | 7.06 | - | 16.21 |
| **L2G4** | MP+MPP | 2.3 | 11.77 | 3.53 | 0.32 | 15.60 |
| **L2G7** |  |  |  | 7.06 | 1.12 | 15.86 |
| **L4G4** |  | 4 | 15.57 | 3.53 | 0.20 | 15.43 |
| **L4G7** |  |  |  | 7.06 | 0.58 | 15.49 |
| **L4G7-RGE** | MP+MPP | 4 | 15.57 | 7.06 | 0.58 | 15.49 |

Traditionally, ligand-functionalised biomaterials only control the global ligand density in a material; however, we and other researchers have demonstrated that local ligand density is also a key yet understudied biomaterial property because it promotes receptor clustering, which is key for driving intracellular signalling ^[15,29,30,33]^. Adherent cells, including myoblasts, primarily engage with the extracellular matrix through integrin receptors that bind peptide ligands in the extracellular space. *β*_1_ and *β*_3_ integrins are primarily responsible for the formation of focal adhesions, which are complex assemblies of transmembrane receptors, adaptor, and signalling proteins that are key signalling hubs, particularly for the transmission of mechanical signals. The integrins must cluster in the cell membrane to form focal adhesions for optimal signalling to occur, and creating interfaces with ligand multivalency is a facile method of achieving this. However, the impact of ligand multivalency on skeletal muscle development and NMJ formation has not been assessed. In this work, we functionalised the polymers with the RGD peptide sequence found in several ECM proteins, such as fibronectin, because it engages multiple integrin receptors (e.g., α_5_β_1_, α_v_β_3_, and α_v_β_5_) necessary for focal adhesion formation^[33–37]^.

Substrate topography is another biomaterial characteristic that can be used to promote myogenesis. Specifically, employing aligned features and fibers with a 1.3-3.0 µm diameter range has been shown to improve myotube formation in C2C12 myoblasts and increase alignment, adhesion, proliferation, and differentiation ^[24,38]^. However, the interaction between ligand multivalency and substrate topography has not been assessed as a method to improve myogenesis and NMJ formation. To fill this research gap, we fabricated the MP+MPP polymer blends into different architectures, including cast films (C), random electrospun fibers (R), and aligned electrospun fibers (A), as shown in **Figure 1A**. Electrospinning was selected as the fabrication method because it is a readily available and facile method of fabricating polymer solutions into substrates with the appropriate micron-scale morphology to promote myogenesis. The polymer fibers were evaluated using SEM (**Figure 1B-E**). SEM images revealed uniform and bead-free fibers with random or aligned fiber morphology (**Figure 1B**). Quantification via ImageJ analysis revealed that the average diameters for random and aligned fibers were 1.87 ± 0.54 µm and 1.78 ± 0.32 µm, respectively, falling within the range previously identified to promote myogenesis ^[38,39]^. The electrospinning parameters were adjusted so that no statistical differences were observed between groups (**Figure 1C**). Additionally, quantification of fiber alignment indicated that aligned samples exhibited directional orientation, with an average directional coherency coefficient of 0.91 (**Figure 1D, E**).

To the authors’ knowledge, peptide nanopatterning on complex, 3D surfaces has not previously been reported; therefore, a variety of surface characterisations were used to compare the presence and distribution of peptides across the biointerfaces, including attenuated total reflection-Fourier transform infrared (ATR-FTIR), energy dispersive X-ray spectroscopy (EDX) spectroscopies, and nanoparticle tagging (**Figure 2**).

**Figure 2.**
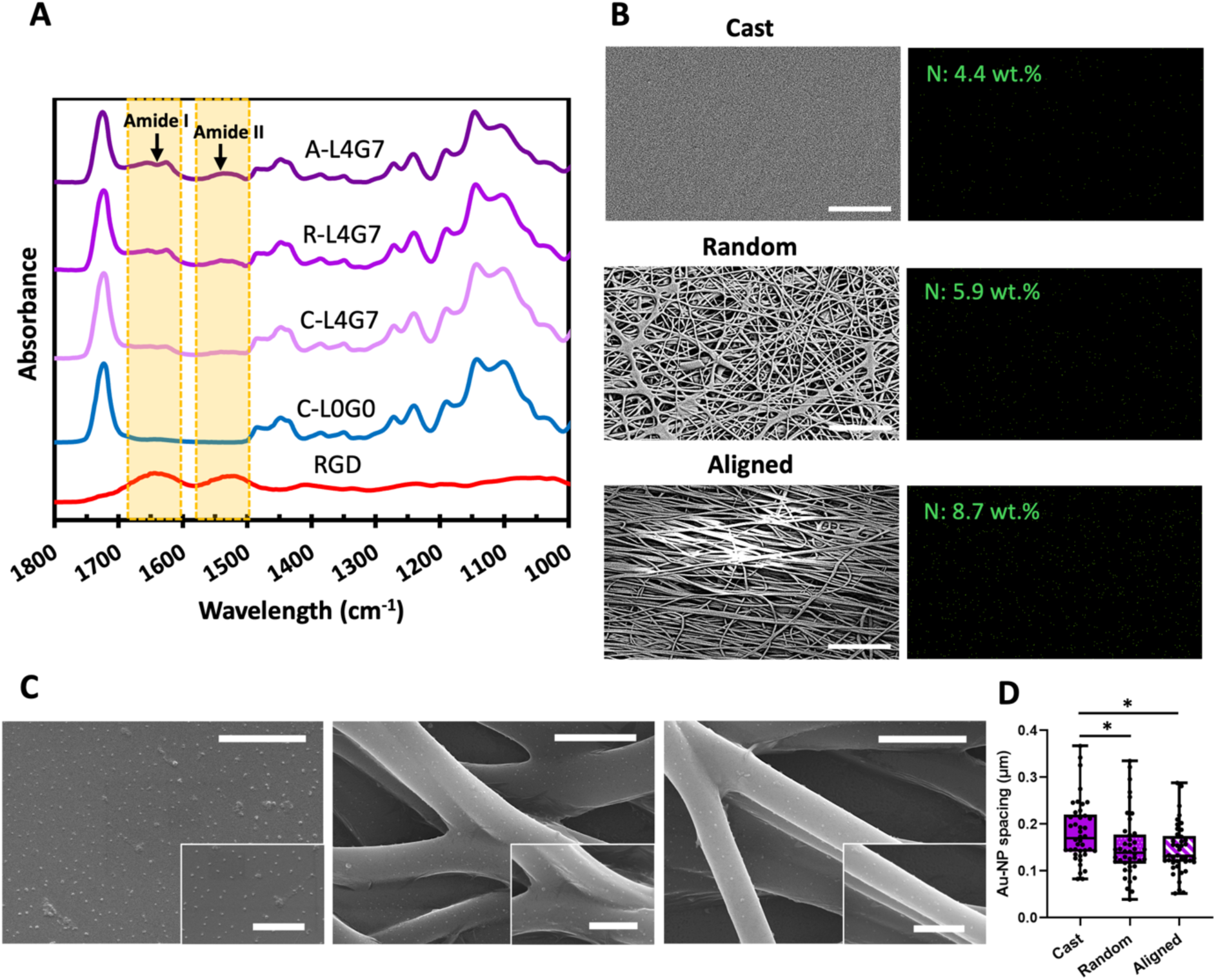
Evaluation of surface composition and peptide distribution. A) ATR-FTIR spectra showed the appearance of the amide peaks, which indicate the presence of peptides at the biointerface. B) EDX mapping confirmed the presence of nitrogen corresponding to amine groups of the peptides at the interface, with an increase in nitrogen content in response to fibrous surface morphology (scale bars: 100 µm). C) SEM images of C-, R-, and A-L4G7 biointerfaces tagged with NHS-AuNPs. D) Image analysis using ImageJ shows the difference in spacing between particles (scale bars are 3 µm and 1 µm for big and small SEM panels, respectively). Asterisks (*) above the bars indicate statistical differences between groups. Statistical significance is indicated as p ≤ 0.05 (*), ≤ 0.01 (**), ≤ 0.001 (***), and ≤ 0.0001 (****). Four measurements per 3 replicates were used for SEM analysis.

ATR-FTIR and EDX mapping demonstrate the presence of peptide at the biointerfaces. The ATR-FTIR spectra shown in **Figure 2A** display characteristic peaks for the unfunctionalized L0G0 polymer, including C-O stretching (1055 cm^-1^) and C=O stretching (1730 cm^-1^), and for pure RGD, including the amide I (1650 cm⁻¹) and amide II (1550 cm⁻¹) peaks resulting from C=O stretching and N-H bending, respectively. The appearance of the amide I and amide II peaks in the L4G7 samples indicates the presence of the RGD peptide near the biointerfaces. EDX mapping results support the ATR-FTIR data (**Figure 2B**). Specifically, EDX mapping also revealed differences in the quantity of nitrogen (uniquely arising from the amide bonds in RGD) at the surfaces. Interestingly, A-L4G7 exhibited the highest nitrogen content compared to R-L4G7 and C-L4G7, with 1.47 and 1.97 times increase, respectively, despite the bulk polymers having a constant peptide density (**Figure 2B**). This observation likely arises due to the samples’ surface morphologies. The fibrous surfaces have a higher surface-to-volume ratio, resulting in a signal from a larger surface area being accounted for in the 2D EDX maps in comparison to the cast films. Additionally, the fibers on the aligned surfaces are more densely packed than the random fiber surface, resulting in more surface area in the 2D EDX map.

We further evaluated the availability and distribution of peptide clusters on the cast and fibrous surfaces by conjugating 10 nm NHS-activated gold nanoparticles (NHS-AuNPs) to primary amine groups present in RGD peptides and analyzing the surfaces with SEM. RGD-containing L4G7 biointerfaces showed robust tagging with the AuNPs (**Figure 2C**) compared to no AuNP tagging of the unfunctionalized L0G0 biointerfaces (**Figure S4**), indicating that the particles were binding to the amine groups present in the peptide. For the image analysis, we assumed that each particle represents one peptide nanocluster (i.e., one MPP polymer chain). Interestingly, there was a ∼1.2-fold decrease in the spacing between the nanoparticles on aligned (∼145.8 nm) and random (∼152 nm) surfaces compared to the cast sample (∼183 nm) (**Figure 2D**), though some of this decrease could be due to the curvature of the fibers, which makes peptide clusters appear closer in 2D renderings. These images demonstrate that the peptide nanoclusters are available for cell binding at the biointerface.

### 2.2. Mechanical properties and topography analysis of polymer samples

The bulk and nanoscale mechanical properties and swelling behavior of the samples were assessed. These parameters are critical because the bulk mechanical properties determine the deformation of tissue engineering scaffolds, and the nanoscale mechanical properties provide the mechanical cues that direct myogenesis ^[40]^. Bulk mechanical properties were measured using a uniaxial tensile tester on samples that were equilibrated in a PBS bath at 37°C (**Figure 3A**). The aligned fibre samples are anisotropic, and the load was either applied in alignment with the fiber direction (A-L4G7) or perpendicular to the fiber direction (AP-L4G7). All samples were loaded until failure, and representative stress-strain curves show significant differences in the samples’ mechanical properties (**Figure 3B**). The Young’s modulus, ultimate tensile strength, and elongation at break for each sample were determined (**Figure 3C-E**). A significant drop in stiffness was observed between the C-L0G0 and the C-L4G7 samples, likely due to increased swelling of the polymer due to the hydrophilic peptide moieties. Additionally, the fibrous scaffolds exhibited significantly different mechanical properties compared to the cast film. Specifically, all fibrous samples were weaker than the cast samples due to their porous structure, which was less able to support load. However, the anisotropic nature of the aligned fibers was demonstrated by the increased strength of the samples deformed in the same direction of fiber alignment (A-L4G7) compared to those deformed in the direction perpendicular to alignment (AP-L4G7). Natural skeletal muscle tissue with a high level of activity has an average tensile modulus in the range of 25-120 kPa ^[41,42]^ and can even reach ∼480 MPa ^[43]^, while the samples assessed in these experiments had similar tensile moduli ranging from approximately 20 – 150 kPa.

**Figure 3.**
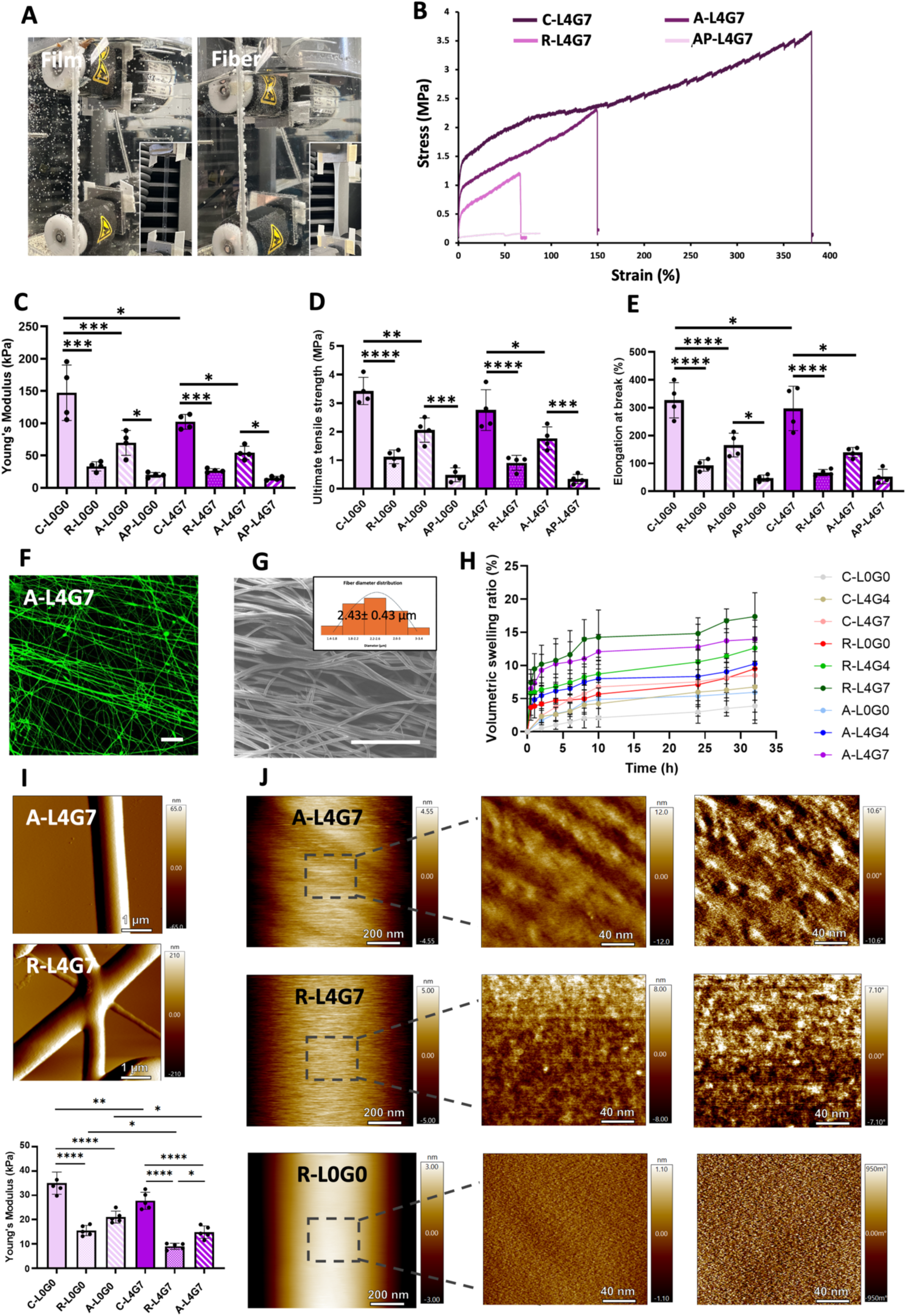
Evaluation of mechanical properties and topography of the polymer scaffolds. A) macroscopic images of samples at approximately 100% strain. B) Representative stress-strain curves for various samples. C) Young’s modulus, D) ultimate tensile strength, and E) elongation at break (n=4). Representative images of aligned fibers (A-L4G7) after 7 days of incubation in PBS at 37°C, including F) confocal microscopy image of FITC-incorporated fibers (scale bar: 50 µm), G) SEM image with fiber diameter distribution histograms (scale bar: 50 µm), and H) quantitative analyses of volumetric swelling ratio of samples (n=5). Representative I) nanoindentation using AFM and calculated stiffness based on JKR model fitting shows variation in stiffness in response to surface morphology and presence of functionalized polymer chains. J) AFM height (left and middle column) and phase (right column) images of polymer fiber samples at equilibrium swelling in water. Four measurements per 3 replicates were used for AFM analysis. Asterisks (*) above the bars indicate statistical differences between groups. Statistical significance is indicated as p-value ≤ 0.05 (*), ≤ 0.01 (**), ≤ 0.001 (***), and ≤ 0.0001 (****).

Significant swelling of the fibrous samples was visualised using confocal images of FITC-incorporated microfibers over 7 days of incubation in an aqueous environment, as seen through an increase in fiber diameter (**Figures 3F** and **S5**). Morphological stability of the fibrous structures was observed after deswelling (**Figure 3G**), where samples retained their aligned or random fiber architecture. The fluorescence micrographs were used to calculate the volumetric swelling ratios (**Figure 3H**), and it was found that swelling increased with peptide density, likely due to the more hydrophilic nature of the peptides.

In addition to bulk mechanical properties, the local, nano-scale mechanical properties are critical design parameters because they are responsible for cellular mechanotransduction that plays a role in directing cellular behavior and function ^[44,45][46,47]^. To evaluate the stiffness of the substrates at microscopic level, atomic force microscopy (AFM) nanoindentation was employed using tapping mode in water on hydrated samples (**Figure 3I**). The average stiffness was measured based on JKR model fitting on the generated force curves, which considers the adhesion effect upon applying the force on soft polymers in water. As with the uniaxial tensile testing, the modulus of the L0G0 group was significantly higher than that of the L4G7 group, for a given surface topography. These results can be explained by increased hydrophilic fraction in the polymer blend, and therefore, enhanced polymer swelling and reduction in stiffness compared to L0G0. Interestingly, the A-L4G7 samples had a statistically higher modulus than the R-L4G7 samples, potentially due to changes in polymer chain orientation during the electrospinning process. The average stiffness values of all indented samples were lower than the modulus from tensile testing, which is likely due to differences in test conditions (compressive vs. tensile force). Accordingly, the measured elastic modulus was 27.75 ± 3.52, 9.02 ± 1.26, and 14.85 ± 2.54 kPa for C-L4G7, R-L4G7, and A-L4G7, respectively.

The polymer fibers were further assessed via AFM height and phase imaging using tapping mode in water to investigate the surface topography single fibers. Topographical mapping demonstrated that fibers without peptide exhibited a very smooth surface (maximum height of ∼1 nm), but a significant increase in surface roughness was observed upon addition of peptide functionalized fraction to the polymer blend (**Figure 3J**). Interestingly, we also observed striations on the surface of the aligned fibers compared with random fibers, potentially due to drawing of the fibre during collecting onto the rapidly rotating collector. Additionally, phase images illustrate the formation of small surface features upon inclusion of the peptide-modified polymer fraction, and these were consistent with our previous findings ^[15]^. This is potentially due to local accumulation of polymer chains containing charged RGD peptides through autophobicity ^[48]^.

### 2.3. Peptide distribution, density, and surface presentation regulate myoblast adhesion, morphology, and proliferation

Fluorescence microscopy was used to determine if C2C12 myoblast cell adhesion and spreading were influenced by surface morphology and peptide density/distribution through staining for actin and vinculin after 24 h of culture. The alignment and spreading of the cells were quantified (**Figure 4A-B** and **S6**). The control samples (L0G0 and RGE) showed minimal cell adhesion, confirming the low fouling properties of the materials and selectivity of the peptides (**Figure S5**). C2C12 myoblasts adhered to all other test surfaces through interactions with adhesive RGD peptides (**Figure 4**). However, the extent of adhesion, cell aspect ratio, and spreading varied significantly with ligand density, patterning, and surface morphology. According to **Figure 4A**, moving from polymer cast film to random and aligned surfaces improved the degree of cellular alignment along the direction of the fibers, with ∼2.5- and ∼1.5-fold increase in alignment scores, respectively. This was also evident from the angular distribution histogram, showing that cells on random and cast film surfaces had a more spread and less elongated morphology. Additionally, as can be seen in **Figure 4A-B**, on peptide cluster surfaces regardless of surface morphology, cells showed increased spreading and stronger actin staining in response to higher local density (approximately 2.5-fold increase in L4G7 compared to L1G7). At this early time point, the A-L4G7 surface appeared best for *in vitro* skeletal muscle development by promoting maximal adhesion and directional alignment of the cells.

**Figure 4.**
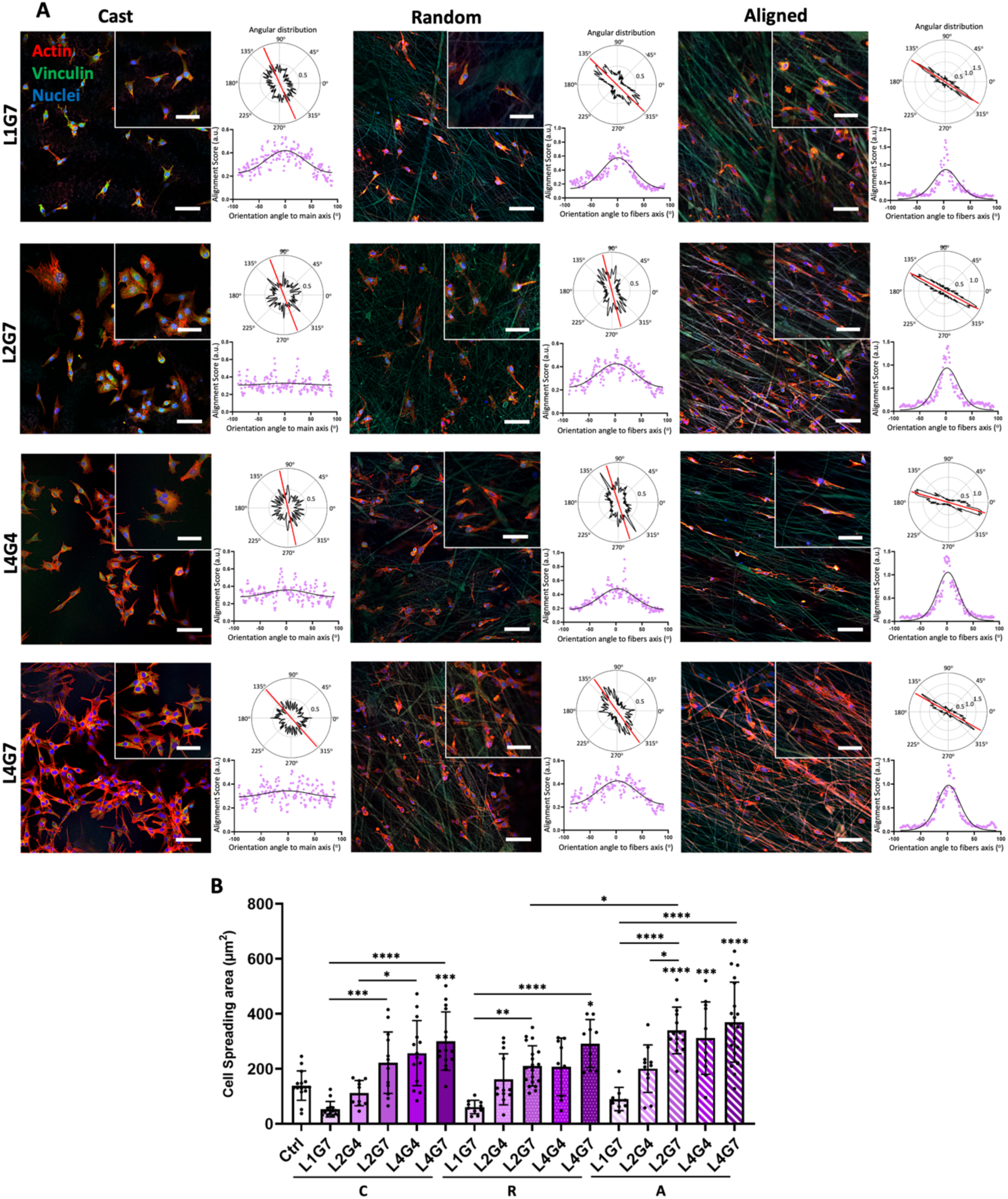
Evaluation of C2C12 myoblast adhesion and orientation in response to ligand density and surface morphology. A) Representative confocal microscopy images of cells cultured on different samples after 24 h with cellular alignment analysis using angular distribution (top) and alignment score with respect to orientation angle (bottom) graphs, and B) quantification of average cell spreading area as a function of ligand density and surface morphology (n=15). Staining shows actin filaments (red), vinculin (green), and nuclei (blue). For better visualization of the cell alignment in the direction of fiber orientation, FITC-labelled fibers were used. Scale bars are 100 µm and 30 µm for the zoomed-in panel. Asterisks (*) directly above data points indicate statistical differences compared to the control, and asterisks above the bars indicate statistical differences between treatment groups. Statistical significance is indicated as p-value ≤ 0.05 (*), ≤ 0.01 (**), ≤ 0.001 (***), and ≤ 0.0001 (****).

**Figure 5** quantifies adherent cell number during 3 days of culture on the test surfaces and shows the morphology and alignment of the cells. The negative control surfaces (L0G0 and RGE) showed minimal cell adhesion over the course of the experiment. In contrast, cells were able to adhere to and grow on all other surfaces, in a substrate-dependent manner. On days 2 and 3, increased ligand density and fiber alignment both played a role in regulating myoblast numbers. Specifically, there was a 1.4-fold increase in cell density on A-L4G7 compared to the positive control on day 3. In addition to having the largest cell population, the A-L4G7 surface also promoted a high degree of cell alignment.

**Figure 5.**
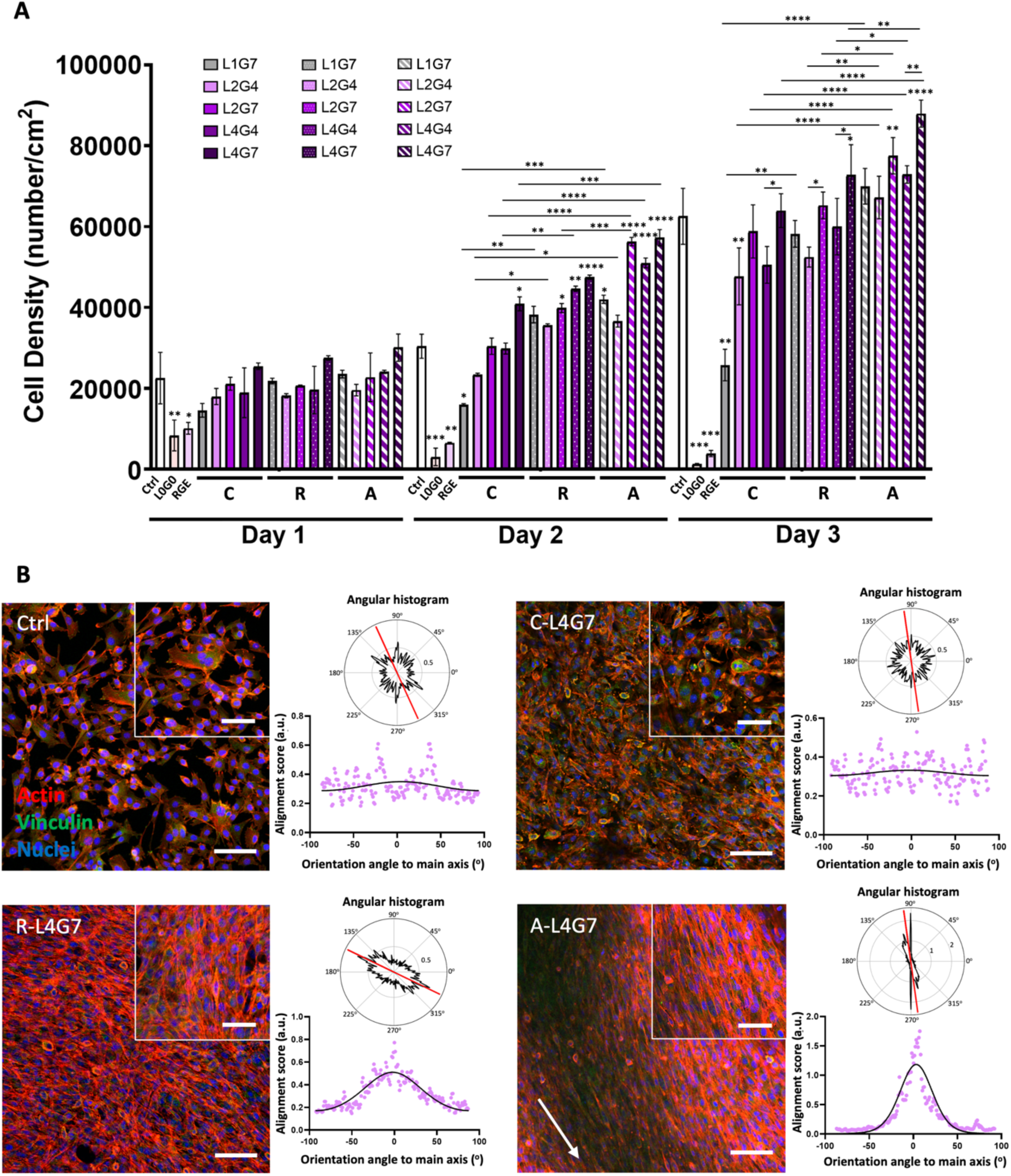
Evaluation of (A) C2C12 myoblast density on different substrates over 3 days. Asterisks (*) directly above data points indicate statistical differences with the control, and asterisks above the bars indicate statistical differences between treatment groups (n=6). Statistical significance is indicated as p-value ≤ 0.05 (*), ≤ 0.01 (**), ≤ 0.001 (***), and ≤ 0.0001 (****). B) Representative confocal microscopy images of cells on highly peptide-functionalized polymer samples (L4G7) with different surface morphologies on day 3, with cellular alignment analysis using ImageJ. Staining shows actin filaments (red), vinculin (green), and nuclei (blue). Scale bars are 100 µm and 30 µm for the big and small panels, respectively. The white arrow shows the direction of fiber alignment.

### 2.4. Myotube formation is regulated by surface morphology and peptide distribution

Proper neuromuscular junction formation *in vitro* requires the differentiation and fusion of myoblasts into myotubes that can connect with motor neurons. Therefore, we assessed the impact of surface morphology and peptide distribution on myotube formation. Myogenic differentiation was evaluated using immunofluorescence staining against two major myogenesis hallmarks: myosin heavy chain (MyHC) and ⍺-sarcomeric actinin. From the resulting confocal fluorescence micrographs, various parameters were quantified, including myotube diameter, aspect ratio, alignment, myoblast fusion index, and sarcomere-forming myotube fraction (**Figure 6** and **S7-8**).

**Figure 6.**
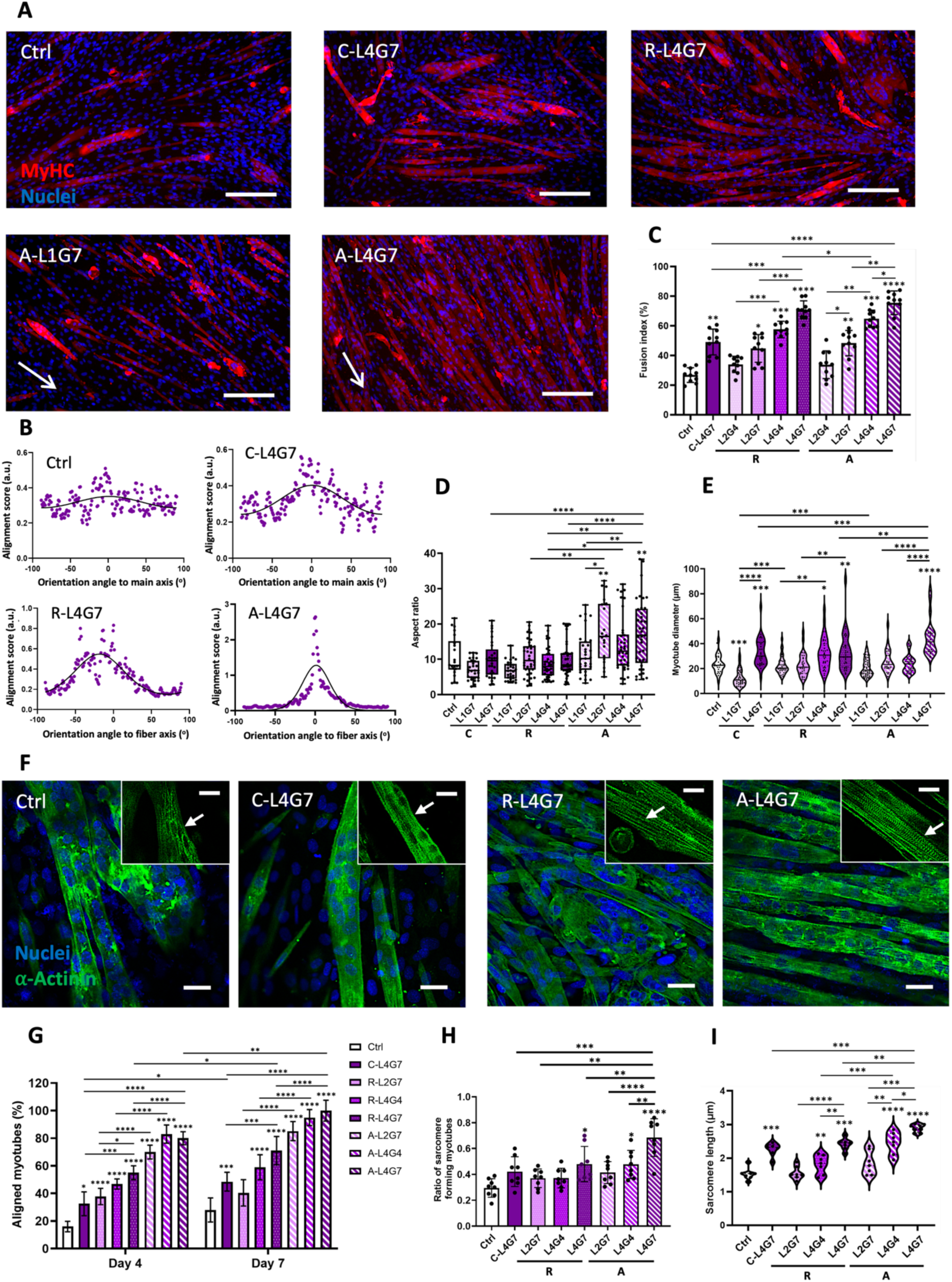
Assessment of myogenesis and maturation after 7 days of differentiation on various biomaterial samples. A) Representative confocal microscopy images of myotubes stained for MyHC (red) and nuclei (blue) (Scale bar is 200 µm and white arrows show the direction of fiber alignment). Cell morphology and myotube development were quantified by B) myotube alignment, C) myoblasts fusion index, D) myotube aspect ratio, and E) myotube diameter using ImageJ (n=6). F) Representative confocal microscopy images against sarcomeric *α*-actinin (green) and nuclei (blue) reveal the formation of z-lines and striation of myofibers, indicating their maturation (indicated by white arrows, scale bar of 50 µm). Images were further assessed to measure G) percentage of aligned myotubes, H) the ratio of sarcomere-forming myotubes to total myotubes, and I) sarcomere length (n=10). Asterisks (*) directly above data points indicate statistical differences with the control, and asterisks above the bars indicate statistical differences between treatment groups. Statistical significance is indicated as p-value ≤ 0.05 (*), ≤ 0.01 (**), ≤ 0.001 (***), and ≤ 0.0001 (****).

Confocal images demonstrated that myotube structure and organization varied across different groups (**Figure 6A** and **S7**). On control and cast film surfaces, myotubes were short and thin and had no apparent alignment. Random fibers promoted the formation of thicker but shorter myotubes with no alignment. In contrast, the aligned surfaces promoted myotube elongation and directional orientation. This directional orientation was further demonstrated by the quantified myotube alignment analysis that showed the sharpest peak for cells on the A-L4G7 substrates (**Figure 6B**). Additionally, surface functionalisation also played a critical role in myogenesis. Both global and local ligand densities increased myogenesis and myotube diameter regardless of surface morphology, with the greatest fusion index achieved on myoblasts differentiated on L4G7 (2-fold diameter increase compared to L1G7) (**Figure 6C-E**).

The cultures were also imaged for α-actinin, a hallmark of myotube maturity and late-stage differentiation (**Figure 6F**). Fluorescence micrographs also demonstrated increased myotube alignment in cultures grown on aligned polymer fibers relative to the random fibers and cast films (**Figure 6F, G**). Furthermore, the micrographs also enabled visualisation of z-lines indicative of sarcomere formation. Quantification of the ratio of z-line-forming myotubes to total myotubes and sarcomere length was performed. The largest fraction of sarcomere-forming myotubes, as well as the longest and most uniform sarcomere length, was found on the A-L4G7 substrates, further demonstrating the importance of ligand density (local and global) and surface alignment for skeletal muscle development *in vitro* (**Figure 6H, I**, and **S8**).

The results illustrated significant improvement in the number, length, and organization of the sarcomeres, particularly on the A-L4G7 surfaces (**Figure 6H, I**, and **S8**). Additionally, the myotubes on these surfaces exhibited spontaneous contraction (**Supporting Information, SIM-1 and 2**), demonstrating a higher degree of maturation and function. While previous studies have shown sarcomere formation in C2C12 cells cultured on aligned scaffolds such as collagen ^[49]^, gelatin ^[50]^, and RGD-functionalized pectin ^[13]^, these studies often required prolonged culture of 2–3 weeks or required external stimulation to induce contraction. The rapid development of organized sarcomeres and spontaneous contractility within just 7 days on peptide-functionalized anisotropic scaffold demonstrates a significant advance in the development of *in vitro* skeletal muscle models.

### 2.5. Formation of neuromuscular junction *in vitro* regulated by surface alignment and integrin binding ligand clustering: impact on AChR clustering and neurogenesis

The successful engineering of a contractile skeletal muscle tissue model necessitates the formation of neuromuscular connections and synapses on myofibers, which is challenging *in vitro* without the application of external electrochemical stimulants *^[1,28,51]^*. To form NMJs, sequential direct co-culturing was used (**Figure 7A-B**). First, myoblasts were differentiated into myotubes, and motor neurons were grown and differentiated to allow lineage commitment and neurite formation in monoculture. Motor neurons were then transferred on top of the myotubes to allow NMJ formation. Neuronal differentiation was confirmed by staining for *β*-tubulin III as a neuronal microtubule marker (**Figure S9**).

**Figure 7.**
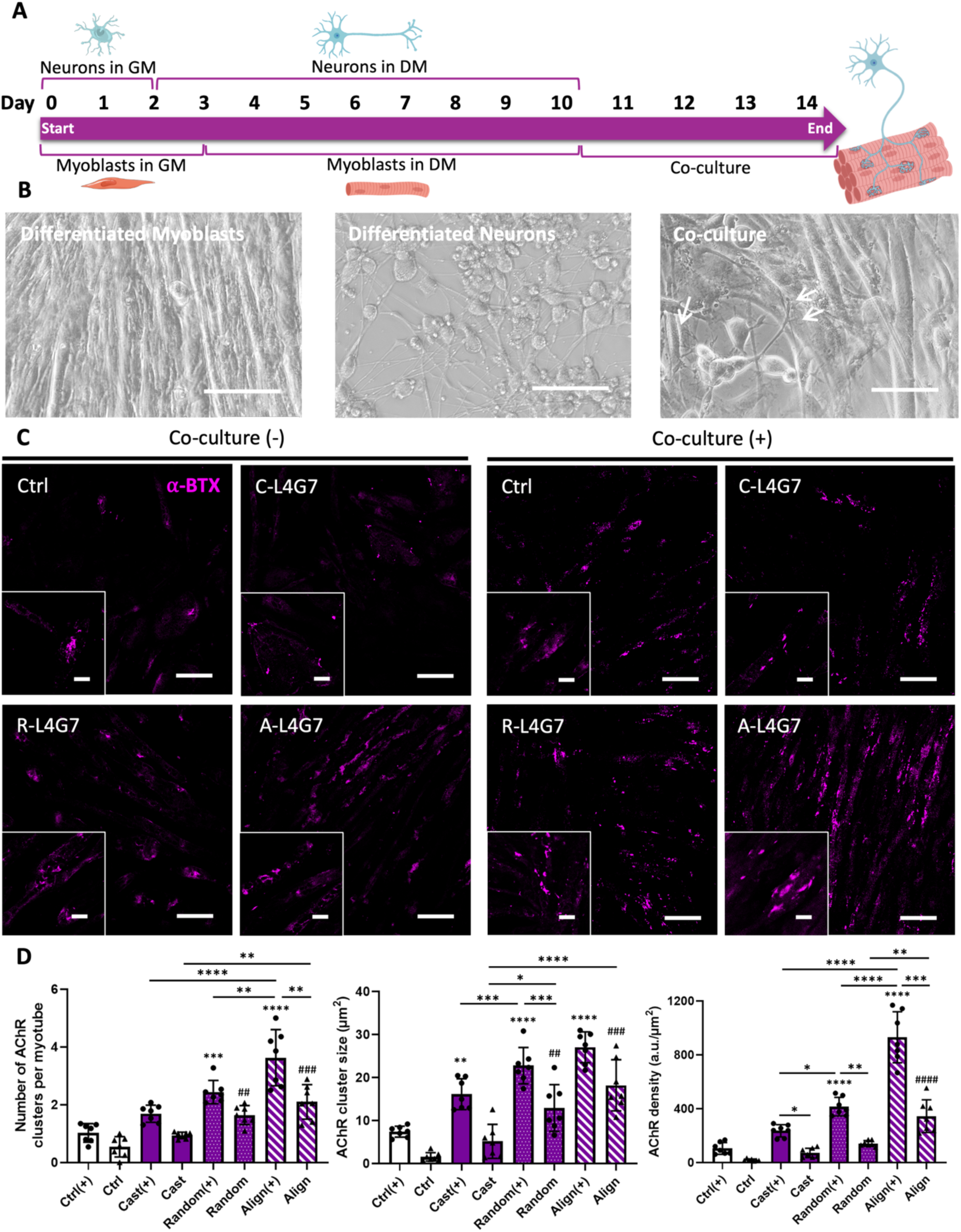
*In vitro* NMJ formation was achieved through A) sequential direct co-culture of myoblasts and motor neurons with B) representative phase-contrast images of differentiated myoblasts, neurons, and their co-culture for NMJ formation. Neural interactions with myotubes are shown by white arrows, indicating the development of NMJs (scale bar: 60 µm). Characterization of NMJ using C) ⍺-bungarotoxin (⍺-BTX) immunostaining (magenta) to visualize AChR under the effect of different surface morphologies and with and without neuronal co-culturing for 4 days (scale bars are 100 µm and 15 µm for images and their zoomed-in version, respectively), and D) quantification of AChR cluster number, size, and density. Asterisks (*) and hashtags (#) directly above data points indicate statistical differences with co-culture control and without co-culture control, respectively. Asterisks above the bars indicate statistical differences between treatment groups (n=10). Statistical significance is indicated as p-value ≤ 0.05 (*), ≤ 0.01 (**), ≤ 0.001 (***), and ≤ 0.0001 (****).

NMJ formations were first demonstrated through immunostaining for α-bungarotoxin (α-BTX), a postsynaptic marker to identify acetylcholine receptors (AChR) clustering on myotube membranes in the presence (co-culture +) or absence (co-culture -) of neuronal co-culture (**Figure 7C**). The fluorescence micrographs were analysed to quantify the number of AChR clusters per myotube, the average size of the AChR clusters, and the density of AChR clusters (**Figure 7D**). Minimal α-BTX staining was observed on surfaces without neuronal co-culture. However, AChR clusters on the A-L4G7 substrates appear more prevalent and larger. Significantly more AChR staining was observed on comparable surfaces with co-culture of motor neurons, with the A-L4G7 surfaces again showing the most staining and larger stained areas. These qualitative results were confirmed via the quantified data, with the A-L4G7 surfaces resulting in the greatest number, size, and density of AChR clusters, and with the presence of motor neurons increasing these parameters compared to myotube monoculture (**Figure 7D**). The increased number and size of AChR clusters with neuronal co-culture is attributed to the ability of neurons to secrete neurotransmitters, including agrin, which plays a crucial role in AChR clustering and NMJ stabilization ^[7,18]^. In this regard, motor neuron-secreted agrin activates the muscle-specific kinase (MuSK) pathway, which drives the aggregation of AChRs at the postsynaptic membrane, thereby increasing the number, size, and density of clusters **Figure 7D**.

In addition to the prevalence and size of acetylcholine receptor clusters, the shape of the clusters is emerging as an important parameter. Specifically, clusters with lacunas indicates postsynaptic organization and NMJ maturation ^[26,52,53]^. Co-cultures of myotube and motor neurons were imaged via confocal microscopy and super resolution microscopy after immunolabeling against MyHC (MF20, red) and AChR (α-BTX, green), and nuclei (DAPI, blue) (**Figure 8A-B**). The number of AChR microclusters per myotube and their lacunarity across the various surfaces were quantified (**Figure 8C-D**).

**Figure 8.**
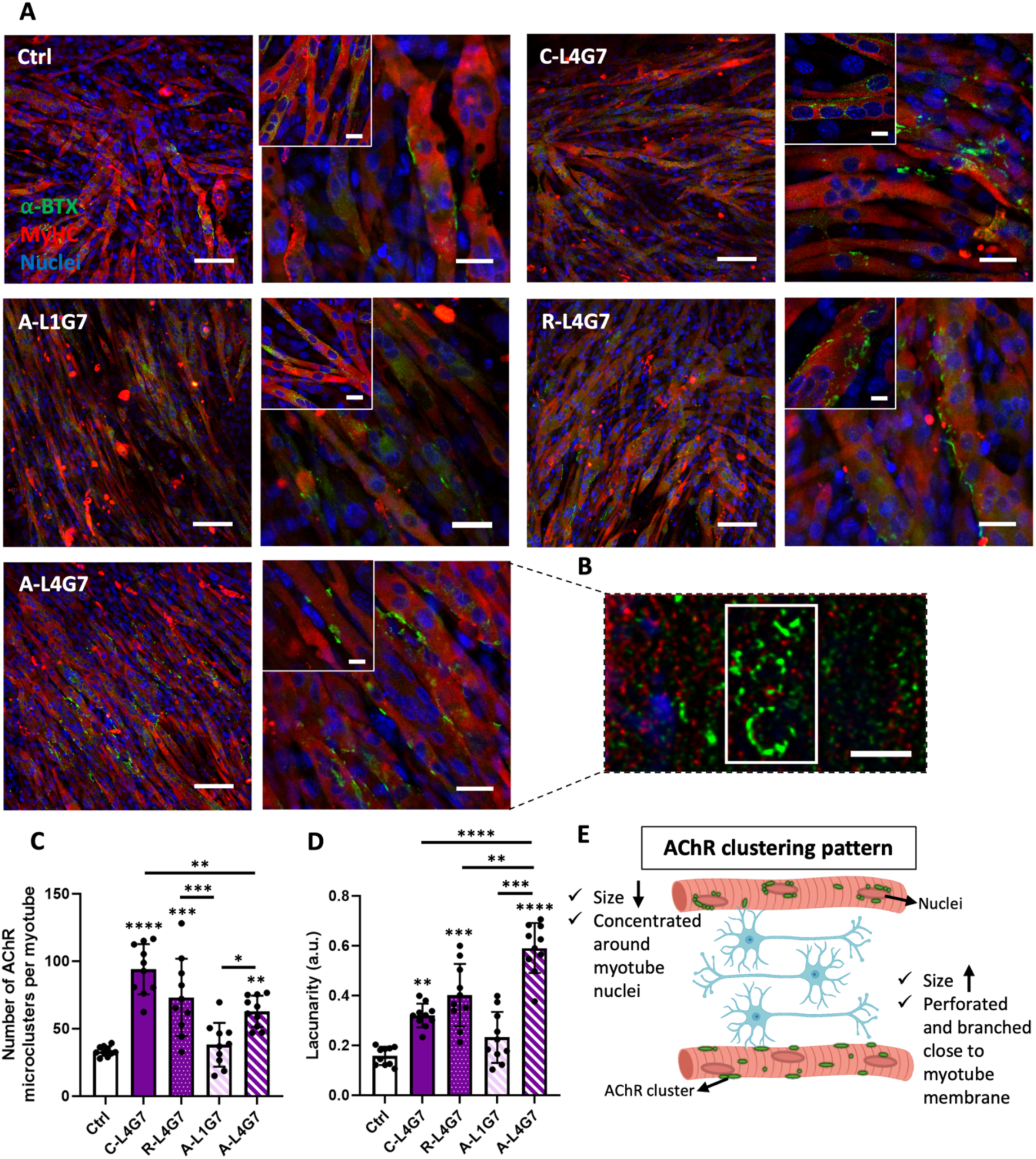
Evaluation of AChR shape and size under the effect of ligand clustering and surface morphology. A) Representative confocal images of myoblast-motor neuron co-cultured samples after 4 days. Formation of perforated AChR clusters and increased number of microclusters were apparent on A-L4G7 and C/R-L4G7 samples, respectively. Staining for MyHC (MF20, red), AChR cluster (⍺-BTX, green), and nucleus (DAPI, blue), scale bar for each sample image: 100 µm (left), 30 µm (right), 15 µm (zoomed-in). B) super-resolution microscopy image of AChR cluster visualizing the branching of the cluster into a C-shape (scale bar: 6 µm). Quantification of cluster size and shapes were presented as C) number of microclusters (size<5 µm) per myotubes and D) lacunarity. Asterisks (*) directly above data points indicate statistical differences with control. Asterisks above the bars indicate statistical differences between treatment groups (n=10). E) schematic summary of two main AChR clustering patterns observed in our study. Statistical significance is indicated as p-value ≤ 0.05 (*), ≤ 0.01 (**), ≤ 0.001 (***), and ≤ 0.0001 (****).

The fluorescence micrographs in **Figure 8A-B** show distinct patterning. AChR clusters on control surface (glass coverslip) and surfaces with randomly distributed ligand (A-L1G7) were small and exhibited low lacunarity suggesting early stages of receptor clustering and less developed NMJs. An increase in cluster number, size, and lacunarity were observed on substrates with multivalent ligands (cast films, random fibers, and aligned fibers). The clusters on cast and random surfaces were bigger with higher lacunarity compared to the control, with the A-L4G7 surfaces showed the highest degree of lacunarity (**Figure 8B**), which is consistent with more mature NMJs. β_1_ integrins play an important role in inducing AChR clustering are regulate aspects of cytoskeletal and membrane organisation in myotubes including the organisation of the postsynaptic apparatus and the presentation of stop signals to growing motor axons ^[18,20,53,54]^. Additionally, integrin binding can result in Rac1 signalling which can mediate membrane remodelling can drive AChR redistribution and cluster stabilization during the later stages of NMJ formation ^[55]^. Thus, the integrin-dependent signaling pathways activated through multivalent RGD binding may regulate the development and maturation of the AChR clusters and NMJ formation.

Previous work has increased the number of AChR clusters on myotubes in co-culture with motor neurons on micropatterned and electrospun fibrous substrates; however, these previous reports usually require electrical stimulation or neurotransmitter treatment to achieve this effect ^[56–59]^, while our biomaterials approach is unique pathway to induce NMJ formation *in vitro* and has significant advantages. Specifically, it does not require expensive and specialised equipment like electrical stimulation bioreactors, and it bypasses challenges associated with neurotransmitter supplementation, including cost and the short half-life of agrin when in culture media, which makes it less effective in the development of postsynaptic machinery ^[18,53,60]^. Additionally, to the best of our knowledge, no other studies except for Bakooshli et al. ^[26]^ have quantified shape factor (lacunarity) as an important parameter for predicting the quality of synaptic organization in cultured myotubes on engineered biomaterials.

We further visualized the neuromuscular junctions with confocal microscopy and SEM (**Figure 9**). For confocal imaging, cultures were stained for motor neurons (β-Tubulin III, yellow), myotubes sarcomeric MyHC (MF20, magenta), and AChRs (α-BTX, green) (**Figure 9A**). The confocal micrographs showed colocalization of neurons, AChR clusters, and myotubes, confirming the formation of NMJs (Pearson’s correlation coefficient of ∼0.7). The arrows highlight regions where the neurites colocalise with myotubes and AChRs cluster, demonstrating synaptic connections. Additional analysis using SEM imaging also confirmed neuromuscular connections (**Figure 9B**). Neurite length and branching were quantified using the confocal micrographs. Although motor neurones on all surfaces exhibited neurite formation; however, a pronounced increase in neurite length (average value of ∼120 µm) and a 1.5-3-fold increase in branches per junction were observed on the aligned surfaces (L4G7) compared to the random (R-L4G7), cast (C-L4G7), and control surfaces (**Figure 9C-D**).

**Figure 9.**
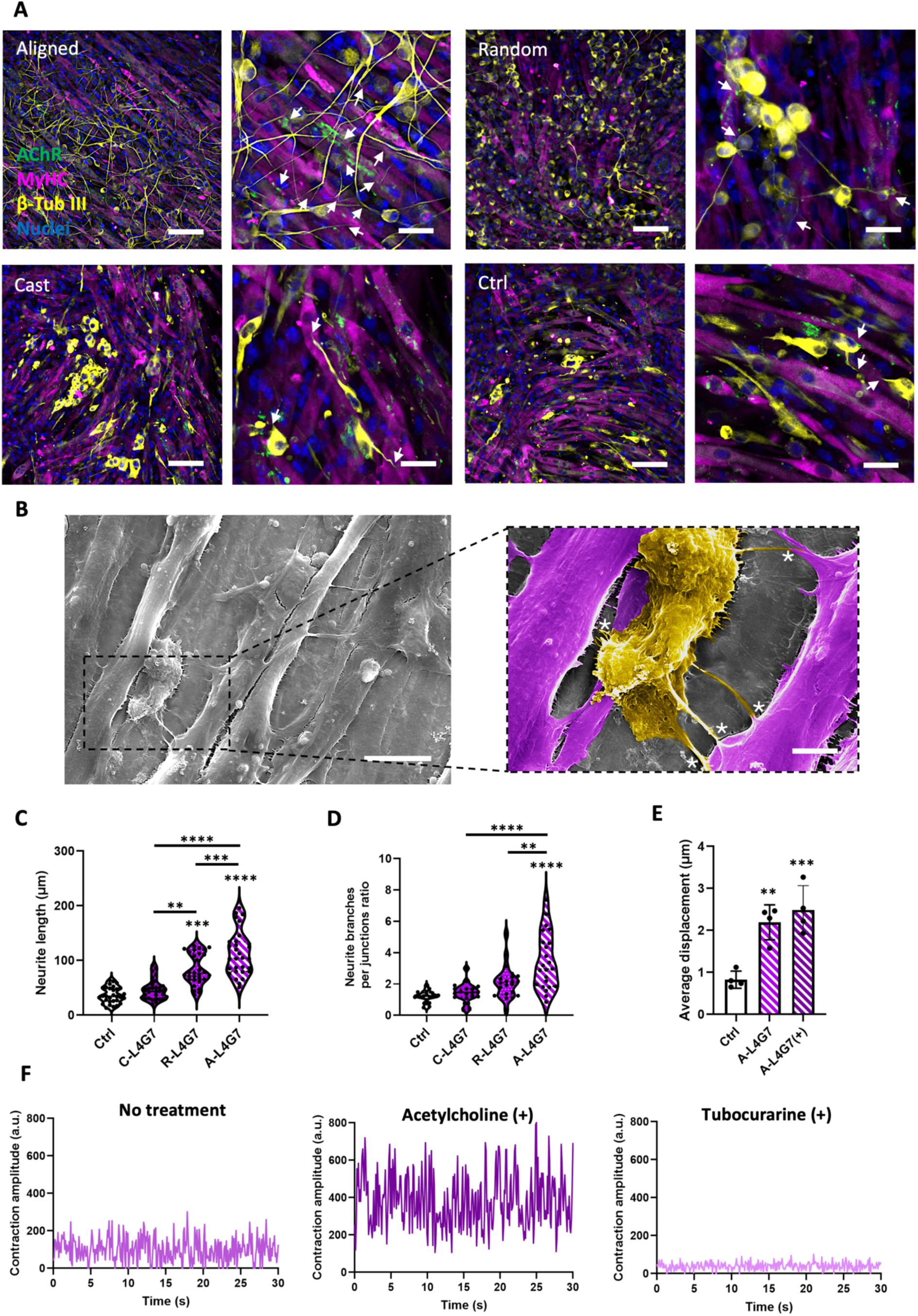
Neuromuscular junction formation and function after 14 days of culture. A) Representative confocal images of neuromuscular co-cultures immunostained for sarcomeric MyHC (MF20, magenta), AChR (⍺-BTX, green), *β*-tubulin III (yellow), and nucleus (DAPI, blue), scale bars: 100 µm (left) and 30 µm (right). Arrows show the AChR clusters co-localized with neurites. B) SEM images of neuromuscular co-cultures further demonstrating motor neuron-myotube connections (myotubes in magenta and neurons in yellow). Scale bars 50 and 10 µm for grey and color images, respectively. White stars (*) indicate neuromuscular connection via extended neurite. C) neurite length and D) neurite branches per junction ratio were determined for the co-cultures as a function of the substrate (n=30). Evaluation of muscle contraction, including E) average displacement of the spontaneous twitching of myotubes before and after co-culture with motor neurons and F) contractile amplitude of myotube twitching with no treatment, stimulated with exogenous acetylcholine, and inhibited with tubocurarine (n=4). Asterisks (*) directly above data points indicate statistical differences with the control, and asterisks above the bars indicate statistical differences between treatment groups. Statistical significance is indicated as p-value ≤ 0.05 (*), ≤ 0.01 (**), ≤ 0.001 (***), and ≤ 0.0001 (****).

The contractile muscle activity at the NMJ was also evaluated after 14 days of culture. We observed spontaneous myotube contraction without exogenous stimulation (**Figure 9F**, **Supporting Information, SIM-1, 2**). Contractions were small and sporadic on control surfaces; myotubes on the A-L4G7 surfaces in monoculture exhibited greater displacement; and the magnitude and synchronisation of the contractions were further increased upon coculture with motor neurons (A-L4G7+) and contractions were observed over the entirety of the surface (**Figure 9F** and **Supporting Information, SIM-3**).

Furthermore, NMJ functionality and cholinergic transmission were further tested by analyzing the muscle contractility after treatment with ACh as an excitatory neurotransmitter to trigger motor neuron firing and contraction, followed by the addition of the ACh antagonist, tubocurarine, which blocks the AChR and disrupts the contraction. As can be seen in **Figure 9G**, treatment of co-culture with ACh enhanced the contraction amplitude of myotubes, and addition of tubocurarine inhibited the contraction (**Supporting Information, SIM-3, 4, 5**).

In the current study, we have developed micro-nano biointerfaces that promote the rapid development of more mature and contractile skeletal muscle *in vitro* as well as their connection to motor neurons in co-culture. This was accomplished by providing cells with nanoscale clusters of integrin-binding peptides and controlling their presentation on anisotropic, fibrous biointerfaces to improve focal adhesion-mediated signaling and structural organization. This approach resulted in accelerated neuromuscular connectivity and robust contractility without an external stimulant. Our results suggest an efficient and promising platform for the facile formation of functional NMJ *in vitro* without the addition of any neurotrophic factor, suitable for neuromuscular disease modelling and drug screening.

## 3. Conclusion

Creating *in vitro* models of skeletal muscle tissue with neuromuscular junctions often requires the provision of complex biochemical and physical cues such as neurotrophic factors or electrical stimulation. In this study, we introduced an engineered biointerface that promotes the rapid development of more mature and spontaneously contractile myotubes with connection to motor neurons. This was accomplished by providing cells with appropriate cues at the nanoscale (multivalency of cell adhesive ligands) and microscale (aligned architecture), demonstrating the need for appropriate and cooperative cues across length scales. The resulting *in vitro* NMJ co-culture system will provide a valuable platform for modelling degenerative neuromuscular diseases, drug screening, and advancing skeletal muscle tissue engineering.

## 4. Experimental Section

### Synthesis and peptide functionalization of low-fouling polymers

Polymers used in this study were synthesized and functionalized according to methods described previously ^[15,29]^. Briefly, low-fouling polymers, including random comb copolymer of methyl methacrylate-polyethylene glycol methacrylate (MMA-PEGMA, named MP) and random comb terpolymer of methyl methacrylate-polyethylene glycol methacrylate-polyethylene glycol methacrylate-norbornene (MMA-PEGMA-PEGMA+Norbornene, named MPP) were synthesized using the RAFT polymerization technique. After the synthesis of cysteine-terminated RGD (CGGGRGDS) and RGE (CGGGRGES) peptides using Fmoc-based solid-phase peptide synthesis, the terpolymer was functionalized with the purified peptides via thiol-ene click chemistry to produce MPP polymers. After synthesis and functionalization, the polymers were dialyzed using MWCO10kDa and MWCO1kDa dialysis membranes, respectively, against MilliQ water for 3 days and then lyophilized and kept at 4°C until required.

### Preparation of polymeric films with RGD ligand nanoclusters and different surface topographies

Blends of functionalized polymer (MPP, with local densities of ∼1, 2, and 4 RGD per chain) and non-functionalized polymer (MP) were prepared using appropriate blending ratios to attain global densities of 4 and 7 µg of RGD per mg of polymer mixture (refer to the **Table 1**) and dissolving the polymers with 10 wt.% in methanol: water (4:1) at 60 °C. Solution of MP and MPP alone with the same concentration and solvent mixture were also used to generate the surfaces without RGD (L0G0) and randomly distributed RGD (L1G7), respectively, as control surfaces along with a polymer blend with the highest global and local densities of control peptide (RGE).

To generate polymeric films, the polymer solution was cast (50 µl.mm^-2^) onto substrates (round glass coverslips, No.1, 11, and 13 mm). For fibrous topography, the polymer solutions were electrospun using an 18G blunt needle with a flow rate of 0.6 ml.h^-1^, under a high voltage of 17 kV, and with a 15 cm distance between the needle tip and the collector. Random and aligned fibers were collected on round glass coverslips or aluminium foil on a rotating mandrel with rotation speeds of 100 and 800 rpm, respectively. Both cast and fibrous samples were slowly dried under a fume hood at room temperature for 24 h before use. For biological assays, all samples were sterilized using UV light for 2 h, 3× rinsing with sterile PBS, and then incubated with serum-free culture media at 37°C overnight before cell seeding.

Different sample groups were referenced based on their topography and ligand content using A, R, and C for Aligned and Random fibers and Cast film, and L and G for local and global densities (as shown in **Table 1**), respectively.

### Scanning electron microscopy (SEM) and Energy dispersive X-ray spectroscopy (EDX)

Surface morphology and presence of nitrogen as indicative of RGD peptides in polymer were evaluated by SEM (Hitachi FlexSEM 1000, Japan) and EDX microanalysis (Bruker, USA) using 20 keV exposure. Vacuum-dried samples were used for SEM and for EDX analysis. After soaking the samples in DI water overnight, they were lyophilized and sputter-coated with gold. Fibers’ average diameter, distribution, and directionality were analysed using DiameterJ and OrientationJ plugins in ImageJ software.

### Attenuated total reflection-Fourier transform infrared (ATR-FTIR) spectroscopy

Samples were lyophilized and further characterized chemically by ATR-FTIR spectroscopy (Bruker LUMOS FTIR Spectrometer, USA) using 32 scans with 4 cm^−1^ resolution to evaluate the covalent peptide conjugation to polymer at the interface.

### Conjugation of NHS-AuNPs

To assess the peptide distribution on the surface, NHS-activated gold nanoparticles conjugation kit (∼10 nm, Cyto Diagnostics, Canada) was used to label the primary amine groups of the peptides on the polymer surface following the manufacturer’s protocol. First, the polymer samples were washed and incubated in MQ water at 37 ℃ temperature overnight before conjugation. The particles were suspended in the provided re-suspension buffer (1 mg.ml^-1^) and sonicated for 1 h at room temperature (RT). Then, samples were incubated in conjugation kit reaction buffer (pH∼8) including suspended particles at 37 °C for 2 h under mild shaking (200 rpm), followed by the addition of quencher solution to stop the reaction. Samples were washed with MQ water and then incubated overnight at 37 °C under mild shaking to remove the unbound particles. Thereafter, samples were dried under vacuum and sputter-coated with a 5 nm layer of platinum for SEM analysis using Hitachi FlexSEM 1000.

### Uniaxial tensile mechanical testing

Macroscopic mechanical characteristics of film and electrospun samples were determined using uniaxial tensile testing (Universal Testing Machine, Instron 5944, US). Samples with an average thickness of 150 μm were cut into 0.5 × 2.5 cm^2^ rectangles (based on^[63]^ and ASTM D882 ^[61]^) and were mounted on the microtester where tension was applied using a 50 N load cell with a strain rate of 1 mm.min^-1^ until 100% strain. All tests were performed in a PBS buffer chamber at 37°C, and Young’s modulus was calculated from the elastic region (n=4). For aligned fibers, the mechanical testing was applied parallel and perpendicular to the alignment direction.

### Volumetric swelling testing

The volumetric swelling of the sample was measured using fluorescence micrographs ^[40]^. To accomplish this, 10 µl from fluorescein isothiocyanate (FITC) (1 mg.ml^-1^) was mixed with the electrospinning solutions (1:100), and fibers were spun following the same spinning parameters. After imaging the dry fluorescent fibrous samples, they were then incubated in PBS at 37 °C for 7 days with shaking to allow the samples to obtain equilibrium swelling. Confocal laser scanning microscopy was used to evaluate volumetric fiber swelling in all groups using 20x objective with a numerical aperture (NA) of 1.2. During the first 36 h of incubation, at specific time points, samples and their dimensional changes were examined. To minimize bias, from each sample with 5 repeats, 5 random locations were chosen, and average fiber diameters in the field of view were measured using ImageJ, and volumetric swelling ratios were analyzed following the equation below. Film samples’ volumetric swelling was also measured for comparison. To further monitor the changes in fiber morphology after 7 days, freeze-dried samples (n=4) were imaged by SEM.

Volumetric swelling ratio 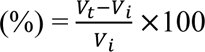

V_i_ and V_t_ are the initial and timepoint volumes of single fibers or film, respectively.

### Atomic force microscopy (AFM) for surface topography and nanoindentation

Fiber surface topography and microscopic mechanical properties of fibers were evaluated by AFM using Asylum Research Cypher in tapping mode in water. Spin-coated polymer films (1000 rpm for 10 s) and electrospun fibers (after 30 min spinning) on silicon wafers were used for the tests. For surface topography, height and phase images of samples after overnight hydration in Milli-Q water were captured using a silicon nitride cantilever with a silicon tip (BL-AC40TS, Asylum Research) compatible with Blue Drive Technology.

Nanoindentation was performed to calculate Young’s modulus of the polymer film or fibers in water using a soft silicon nitride cantilever with a silicon nitride tip (MLCT, Bruker, cone shape tip with 17 nm radius) and a spring constant of 45 N.m^-1^. The cantilever’s spring constant was calibrated by determining the InvOLS (Inverse Optical Lever Sensitivity) parameter and subsequent thermal adjustment after indenting on a clean silicon wafer. Young’s modulus of the samples was calculated from individual force curves of five distinct places on each surface (three replicates), which were analyzed and fitted to the JKR (Johnson-Kendall-Roberts) model using the Asylum program.

### C2C12 myoblast culture

Murine myoblasts (C2C12s, ATCC, Manassas, VA, USA) were grown in growth media (GM) containing Dulbecco’s Modified Eagle Medium-High Glucose (DMEM-Glutamax, Gibco, Invitrogen) supplemented with 10% v/v Fetal Bovine Serum (FBS, Gibco, Invitrogen) and 100 units.ml^-1^ penicillin and streptomycin (P/S, Gibco, Invitrogen) and incubated at 37 °C in a humidified, 5% CO_2_ atmosphere. The culture medium was changed every 2 days, and cells at 70% confluency (passage number <10) were used for seeding.

### Myoblast cell morphology and proliferation

C2C12 myoblasts were seeded on polymer-coated or electrospun onto coverslips in ultra-low attachment 24-well plates (Corning) with 2 × 10^4^ cells per well. After 24 h, cell adhesion and morphology were evaluated by immunostaining. Briefly, PBS-washed samples were fixed in 4% paraformaldehyde for 20 min, washed 3 times with PBST (PBS + 0.3% Triton X-100 (Sigma)), and blocked for 1 h in 4% BSA solution in PBST. Samples were then incubated in a solution of rabbit polyclonal anti-vinculin (1:250, PA5-29688, Invitrogen) diluted in PBST and 2% BSA at 4 °C overnight. After 3 × 10 min PBS wash, samples were incubated in a solution of anti-rabbit Alexa Fluor 488 (1:500, A21206, Invitrogen) for 2 h followed by 1 h incubation in ActinRed 555 ReadyProbes (R37112), and 10 min in DAPI (1:1000, D3571) (Life Technologies) and final wash with PBS for 4 times. Samples were imaged using a Nikon A1R+ confocal laser scanning microscope (20x magnification and NA of 1.2). The cell spreading area and orientation were analyzed with ImageJ ^[62]^.

The alamarBlue assay was employed to assess cell proliferation during 3 days of culture. Briefly, 1 × 10^4^ cells per well of 24-well plates were seeded on samples, and each day, GM was removed and replaced by fresh GM containing a 10% v/v solution of 140 µg.ml^-1^ resazurin in PBS (Sigma) and incubated for 3 h. The fluorescence intensity of each well was recorded using a TECAN infinite M200PRO microplate reader (Ex/Em wavelengths: 530/590 nm). The fluorescence intensity was then converted to cell density using a calibration curve generated from absorbance data of multiple cell densities.

### Myoblast differentiation into myotubes

C2C12 myoblasts were seeded on polymer-coated or electrospun onto coverslips at a density of 1.5 × 10^4^ cells.cm^-2^. After 3 days incubation in GM, once confluent, the media were changed to differentiation medium (DM) containing DMEM-Glutamax supplemented with 2% v/v horse serum (HS, Gibco, Invitrogen) and 1% v/v P/S and maintained for differentiation into myotubes for 7 days (media changes every 2 days). At days 4 and 7 of differentiation, cells were stained following the previously described procedure using overnight incubation in solutions of MYH1E Antibody (MF20)-DSHB (uiowa.edu) and α-actinin primary antibodies (EA-53, MFCD00164521, Sigma) at 1:100 and 1:250 dilutions, respectively, and subsequent incubation in secondary antibodies solution of anti-mouse Alexa Fluor 555 (1:500, A-21422, Invitrogen) and anti-mouse Alexa Fluor 488 (1:500, A-11001, Invitrogen) and DAPI (1:1000) for 2 h before imaging with the Nikon A1R+ confocal laser scanning microscope (20x and 60x magnifications, NA of 1.2). ImageJ analysis was employed to measure fusion index, myotube diameters, orientation, aspect ratio, and the ratio of sarcomere-forming myotubes. Analysis of sarcomere length and uniformity was conducted using SotaTool software.

### Motor neuron in vitro culture and NMJ formation

Murine motor neurons (NSC-34, ATCC, Manassas, VA, USA) were maintained in GM (DMEM-Glutamax supplemented with 10% v/v FBS, and 100 units.ml^-1^ P/S) and incubated at 37 °C and 5% CO_2_ atmosphere. The culture medium was changed every 3 days, and cells at 90% confluency (passage numbers 8-10) were used for seeding. For neuronal differentiation, after 2 days, the GM was changed to DM (Neurobasal medium (NBM, Gibco, Invitrogen) supplemented with 2% v/v B-27 (Gibco, Invitrogen), 0.25% v/v L-glutamine (Invitrogen), 1 µM retinoic acid (Abcam), and 100 units.ml^-1^ P/S) and differentiated for 7 days for the sequential direct co-culture based on the timeline shown in (**Figure 9A**).To form neuromuscular junctions, differentiated neurons were gently removed from the culture flask using PBS diluted trypsin (Gibco, Invitrogen) with a 1:0.5 ratio and seeded with a density of 1 × 10^4^ cells/cm^2^ on top of the grown myotubes on the samples for another 4 days. After one day of incubation in a 1:1 mixture of neuronal and myotube DMs, the co-culture samples were kept in neuronal DM supplemented with 1% v/v HS until the end of the experiment.

### Evaluation of NMJ using immunostaining

The average number, size, and density of AChR clusters in co-culture samples were visualized and quantified using immunostaining to detect the expression of α-bungarotoxin (α-BTX) (based on the protocol in ^[26]^) and compared with samples without neuronal co-culture. To label AChRs, the previously mentioned immunostaining process was applied using α-bungarotoxin-Alexa Fluor 488 (1:500, Invitrogen) in a secondary antibody solution and incubated for 2 h. The neurites of differentiated neurons were also immunostained with an antibody solution of mouse monoclonal anti-*β* tubulin III (1:2000, G7121, Promega) and subsequently stained with anti-mouse Alexa Fluor 647 (1:500, A-21235, Invitrogen). The colocalization of immunolabeled myotubes/neurons and AChR was conducted by the JACoP plugin of ImageJ, in which a Pearson’s correlation coefficient of higher than 0.5 was considered as perfectly overlapped fluorescent signals.

Images were captured using the Nikon A1R+ confocal laser scanning microscope and at 20x and 60x (oil immersion lens) magnifications (NA: 1.2), from at least 5 random positions on each sample. The ImageJ particle analyser was used to identify AChR cluster outlines (with binarizing the cluster outlines), and clusters less than 4.5 μm² (microclusters) were removed from further analysis. Cluster numbers were calculated by counting AChR clusters in each image close to neurites and normalising them to the number of MyHC-positive myotubes. The size and density of AChR clusters were measured and averaged across groups using ImageJ ROI and fluorescent intensity analysis. Additionally, to evaluate morphological heterogeneity between AChR clusters on different surfaces, fractal analysis of α-BTX-stained samples was performed using the FracLac plugin in ImageJ ^[26]^. Accordingly, the AChR clusters outline was binarized and lacunarity was calculated and averaged for each experiment. The Super-resolution microscope Elyra 7 (Zeiss with Lattice SIM^2^) with 60x magnification (NA of 1.4) was also employed to visualize the clustering of the AChR within the myotubes’ membrane.

The spontaneous contractility of myotubes was recorded using a Nikon D700 camera (30 fps) connected to an inverted phase contrast microscope (Olympus CKX41, 20x and 10x magnifications, NA of 1.1). For testing NMJ functionality and cholinergic transmission based on ^[28]^, the contraction was recorded before any treatment, then after the addition of acetylcholine (Sigma Aldrich, 10 μM) as an excitatory neurotransmitter, and finally after the addition of tubocurarine chloride hydrochloride pentahydrate to block the contraction (Abcam, 10 μM). The videos (1 min, 3 repeats from 4 different spots) were analysed by the MuscleMotion plugin in ImageJ software.

At the end of co-culture, samples were fixed with 2.5% glutaraldehyde, gradually dehydrated using serial ethanol solutions, and gently dried under vacuum. Gold sputter-coated samples were imaged using the SEM Hitachi FlexSEM 1000.

### Statistical analysis

Data in this study are reported as mean ± standard deviation, and using GraphPad Prism 9 software, they were graphed and statistically analyzed through one-way and two-way ANOVA, followed by the Tukey-Kramer multiple comparison test (significance level of p-value < 0.05). Unless indicated otherwise, all tests were carried out with at least three independent experiments and technical repeats (n). Statistical significance is indicated as p ≤ 0.05 (*), ≤ 0.01 (**), ≤ 0.001 (***), and ≤ 0.0001 (****).

## Acknowledgements

This work was partially supported by the Australian Research Council (FT190100280). S.N. gratefully acknowledges the support of the University of Melbourne via the Melbourne Graduate Research Scholarship. S.N. would like to thank Dr Negar Mahmoudi and Dr Hao Tran for providing insight about working with neurons, and Ms Miao Chen and Ms Nafiseh Olov for assisting with contraction video capture. Also, the authors would like to appreciate the Materials Characterization and Fabrication Platform (MCFP) (especially Dr. Alejandra Ramirez Munoz, for her assistance with super-resolution microscopy) at the University of Melbourne and the Protein Characterization and Peptide Synthesis Platform at Bio21 Institute (Dr. Troy Attard).

## Data Availability

Data will be made available on request.

## Supporting Information

Supporting information is available from the author.

Table of Content

This study investigates the cooperative effects of adhesive ligand distribution, density, and biomaterial morphology for innervated skeletal muscle tissue engineering. Ligand nanoclusters were created in aligned micropatterns by electrospinning of a RAFT-synthesized polymer with controlled ligand presentation. Improved myogenesis, myotubes alignment, neuromuscular connections with enhanced acetylcholine receptor clustering, and contractility highlight the potential of designed biomaterial for neuromuscular disease modelling.

**Engineering *in vitro* models of skeletal muscle with neuromuscular junctions using hierarchical micro-nano biomaterials: Cooperative effect of adhesion ligand nanoclustering and surface anisotropy**

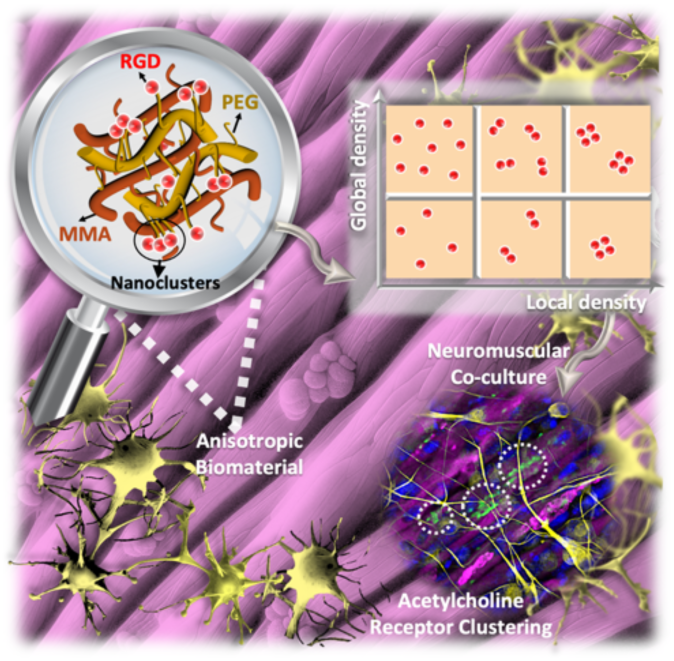

