## Supporting Information for "Engineering *in vitro* models of skeletal muscle with neuromuscular junctions using hierarchical micro-nano biomaterials: Cooperative effect of adhesion ligand nanoclustering and surface anisotropy"

### 1. Experimental Section

#### 1.1. Materials

Methyl methacrylate (MMA, 99%), Poly(ethylene glycol) methyl ether methacrylate (PEGMA, Mw: 475 g.mol$^{-1}$), PEGMA-OH ($M_w$ = 668 g.mol$^{-1}$) synthesized monomer, 2,2′-Azobis(2-methylpropionitrile) (AIBN, RAFT initiator, 98%, ACROS ORGANICS), triethylamine (TEA), 4-Dimethylaminopyridine (DMAP), 5-Norbornene-2-carboxylic acid (mixture of endo and exo, predominantly endo), N-(3-Dimethylaminopropyl)-N′-ethylcarbodiimide hydrochloride (EDC-HCl, Combi Block), 2-phenyl-2-propyl benzodithioate (99% RAFT agent), tris(2-carboxyethyl) phosphine, 2,2-dimethoxy-2-phenylacetophenone (DMPA, photoinitiator), tris(2-carboxyethyl)phosphine, trifluoroacetic acid (TFA), triisopropylsilane (TIPS), N, N′-diisopropylcarbodiimide (DIC) and solvents were obtained from Sigma-Aldrich (Australia) and utilized without any purifying procedures unless otherwise specified.

Mimotopes Pty Ltd (Australia) supplied the amino acids, rink amide resin, and oxyma for the peptide synthesis. To remove inhibitors, the MMA and PEGMA were passed through basic alumina columns and kept at temperatures ranging from 2 to 8 °C. The hydroxyl-terminated poly (ethylene glycol) methacrylate (PEGMA-OH, Mw = 668 g/mol) monomer was synthesized using the method described in [1].

### 1.2. Methods and Results

#### *1.2.1. Polymer synthesis using the RAFT method*

Polymers were synthesized based on the previously mentioned protocol [1,2] via the thermal RAFT polymerization technique. The molar ratio of 1:0.2:1000 for RAFT agent: AIBN: monomers with MMA/PEGMA (M ratio: 850/150) for unfunctionalized polymer (MP) and MMA/PEGMA/hydroxyl-terminated PEGMA (M ratio: 850/50/100) for functionalized fraction (MPP) was used.

After dissolving in 2.5 M dimethylformamide (DMF), the mixture was frozen-pump-thawed three times to remove oxygen, and the reaction flask was filled with argon gas. The polymerization was started by immersing the flask in a 70 °C oil bath with continual stirring for 19 h. The reaction was then stopped by cooling the flask to ambient temperature and opening it to the air. The resultant polymers were precipitated in cold diethyl ether (DEE) and centrifuged three times at 3000 rpm for 5 minutes each to eliminate unreacted compounds.

The vacuum-dried polymer was kept at 4°C until needed. The RAFT agent end groups (i.e., thiocarbonylthio) were then eliminated by mixing the polymer with EPHP at a weight ratio of 50:1 in DMF (total concentration of 102 mg.ml-1). After 30 min of nitrogen bubbling, the mixture was exposed to UV radiation (λ=365 nm) for 24 hours while being stirred continuously. After that, the polymer was dialyzed against water for two days using a dialysis bag with a MWCO of 10kDa, and the resultant polymer was lyophilized and kept at (4°C). The composition of MP and MPP polymers, monomer conversion, and molecular weight distribution were determined based on $^1$H NMR and DMF-GPC (**Figure S1, 2**).

#### *1.2.2. Norbornene functionalization of MPP polymer via an esterification reaction*

Esterification with norbornene (NB) groups was carried out in order to functionalize the pendant OH groups of the MPP polymer with peptides. Briefly, in dry DCM (4 ml) at room temperature under argon for 24 hours, 5-norbornene-2-carboxylic acid (0.059 g, 0.427 mmol) was added to a mixture of polymer (0.1 g), TEA (0.059 ml, 0.423 mmol), and EDC-HCl (0.082 g, 0.428 mmol). Following two centrifugations and precipitation in cold DEE to remove unreacted species from the resulting material, the precipitate was dissolved in DCM and precipitated into distilled water. The polymer was then lyophilized after being dialyzed for two days against a 10% aqueous methanol solution in a dialysis bag with a MWCO of 1 kDa. DMF-GPC and $^{1}H$ NMR were used to examine the resulting polymers (**Figure S2**).

#### *1.2.3. Solid phase peptide synthesis and peptide conjugation using thiol-ene click reaction*

The integrin-binding peptide (CGGGRGDS) and its control (CGGGRGES) were synthesized on Rink amide resin using the Fmoc-based solid-phase peptide synthesis technique for amide C-terminated peptides (resin loading: 0.649). Cysteine amino acids are added into the N-terminus sequence of peptides, together with extra glycine amino acids, to facilitate peptide attachment to NB-functionalized polymer via the thiol-ene process, allowing physicochemical flexibility for peptide sequences in contact with cells. The amino acid to resin coupling was performed using the CEM Liberty Blue microwave peptide synthesiser and auto calculation.

Using a 4- or 5-fold molar excess of Fmoc-protected amino acid (0.4 or 0.5 mmol) that was activated with 0.5 M DIC and 1 M Oxyma, respectively, all peptides were synthesized on a 0.1 mmol scale. Fmoc deprotection was performed with 20% v/v piperidine in DMF. Following a 10- to 15-minute DCM and DMF wash, the resin-containing peptides were vacuum-dried. A cleavage cocktail consisting of TFA, TIPS, and water (92.5:2.5:5, 15 ml/0.1 mmol of peptide) was then used to cleave the peptides off the resin over the course of two hours while being gently shaken continuously.

After filtering and evaporating the peptide-containing cleavage cocktail under nitrogen, the residue was precipitated using ice-cold diethyl ether and centrifuged for three to five minutes at 3000 RPM. Pellets were centrifuged, dried under nitrogen, and washed three times with ice-cold diethyl ether. LC Shimadzu Prep HPLC and a C18 Phenomenex 21.2 mm ID preparative column were used to purify the peptides. A C18 Phenomenex 4.6 mm ID analytical column (5 gradient mode column with buffer A; 0.1% aq. TFA and buffer B; 0.1% TFA in acetonitrile) was used to assess the mass profile and purity of lyophilized peptides. Thermo OrbiTrap Exactive Mass Spectrophotometer and buffer B (0–40% is performed over 40 min, with UV monitoring) are used for gradient elution, respectively. The purified peptides were stored at -20 °C (**Figure S3**).

Thiol-ene click chemistry was used to attach cysteine-flanked peptides to the norbornene functional groups in the polymer because they contain a thiol group at the beginning. 5 mol% of the peptide was combined with 50 mg of MPP-NB, 1 mg of DMPA, and 0.1 mol% of tris(2-carboxyethyl) phosphine in 2 ml of DMF to obtain a 5 mol% norbornene functional group. The mixture was dialyzed (MWCO = 1kDa) against deionized water for 24 hours following 10 hours of UV irradiation. The peptide-functionalized polymers (MPP-RGD) that were produced were freeze-dried and stored at 4°C. Trace elemental microanalysis in CHNS mode was utilized to ascertain the nitrogen content in the polymer and, consequently, the peptide densities in bulk and per polymer chain, utilizing the equations in [1] (**equations 1 and 2**) after the peptides' thiol-ene click reaction was first confirmed by $^{1}$H NMR.

### 2. Results

#### 2.1. Polymer characterizations with NMR and GPC

To determine the composition of the monomer and polymers in deuterated chloroform, proton nuclear magnetic resonance ($^{1}$H NMR) spectroscopy was performed using a Varian Unity Plus 500 MHz NMR spectrometer (CDCl3, 99.8% purity from Cambridge Isotope Laboratories). GPC analysis was performed using a Shimadzu liquid chromatography system and three Phenomenex Phenogel columns (with porosities of 500, 104, and 106 Å; bead size: 5 mm). The columns were kept at 45 ± 1°C. An interferometric refractometer, the Wyatt OPTILAB DSP (functioning at 633 nm), was used. The mobile phase was dimethylformamide (DMF) of high-performance liquid chromatography (HPLC) quality, which flowed at a rate of 1 ml/min. The number-average molecular weight (Mn), weight-average molecular weight (Mw), and polydispersity index (PDI) of the polymers were calculated by calibrating the system with tightly dispersed poly(methyl methacrylate) standards.

**Figures S1 and 2** show the $^{1}$H NMR and GPC characteristics of synthesised polymers in deuterated chloroform and DMF, respectively. Following the polymer purification procedure, the molar ratio of MMA to PEGMA was established by analyzing their unique peaks in the $^{1}$H NMR spectra (see equations in **Table S1**).

As evident in **Table S1**, RAFT polymerization, as a simple controlled living radical polymerization, generated polymers with a high molecular weight (128 and 191 kDa) and low PDI (~1.1), enabling approximate control over the size of the polymer random coil and spacing between peptide-functionalized chains. In addition to $^1$H NMR, the density of peptides in the bulk polymer was calculated by trace elemental microanalysis (**Table 1**).

In MP equations, $X_{PEGMA}$ and $X_{MMA}$ represent the mole percentages of PEGMA and MMA residues in the polymer, respectively. $I_C$ is the integral area corresponding to the $CH_2$ peaks of the $COOCH_2$ methylene group protons inside the repeating unit of PEGMA, whereas $I_A$ is the integral area associated with the $CH_3$ peak for the $CH_3$ methyl group protons in the backbone of both MMA and PEGMA.

In MPP calculations, $X_{PEGMA+PEGMAOH}$ and $X_{MMA}$ represent the mole percentages of PEGMA+PEGMAOH and MMA residues in the polymer, respectively. $I_C$ is the integral area associated with the $CH_2$ peaks that correspond to the $COOCH_2$ methylene group protons inside the repeating units of PEGMA+PEGMAOH. In contrast, $I_A$ is the integral area of the $CH_3$ peak corresponding to the $CH_3$ methyl group protons in the backbone of MMA, PEGMA, and PEGMAOH. Furthermore, $X_{PEGMA}$ and $X_{PEGMAOH}$ represent the mole percentages of PEGMA and PEGMAOH residues in the polymer, respectively. $I_F$ is the integral area associated with the $CH_3$ peaks that correspond to the $CH_3$ methyl group protons inside PEGMA's repeating units.

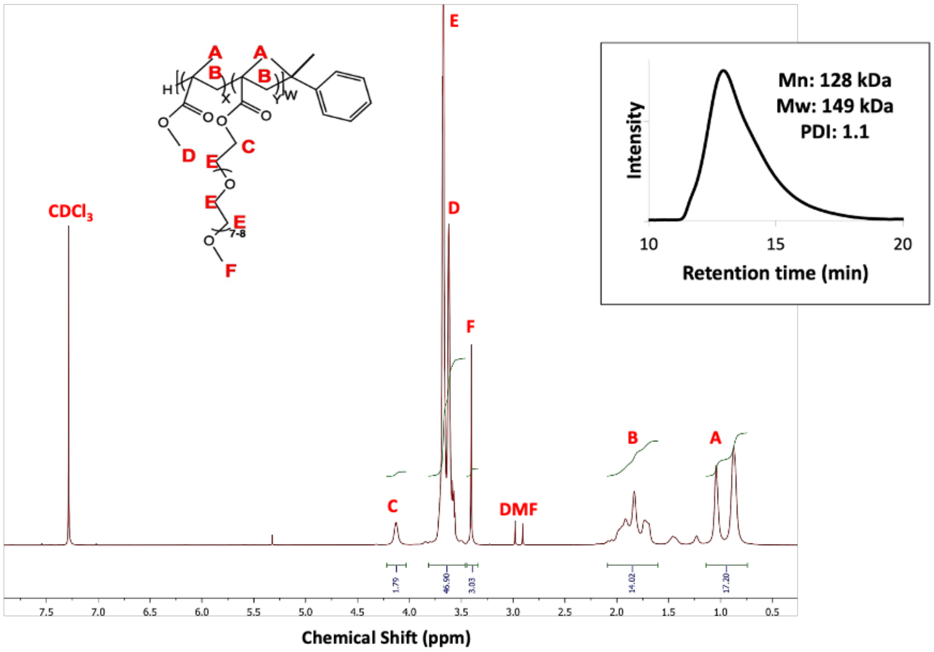


**Figure S1.** $^{1}$H NMR spectrum in deuterated chloroform of MP polymer with GPC spectrum in DMF.

Based on **Figure S1**, for MP polymer, $^{1}$H NMR ($CDCl_3$) includes δ 4.15 (s, 3H), δ 3.62-3.8 (m, 3H), δ 3.5-3.62 (m, 1H), δ 3.4 (s, 1H), δ 1.6-2.1 (m, 1H), δ 0.68-1.12 (d, 1H).

### 2.2. Evaluation of norbornene functionalization of polymers and conjugation of synthesized peptides to polymer

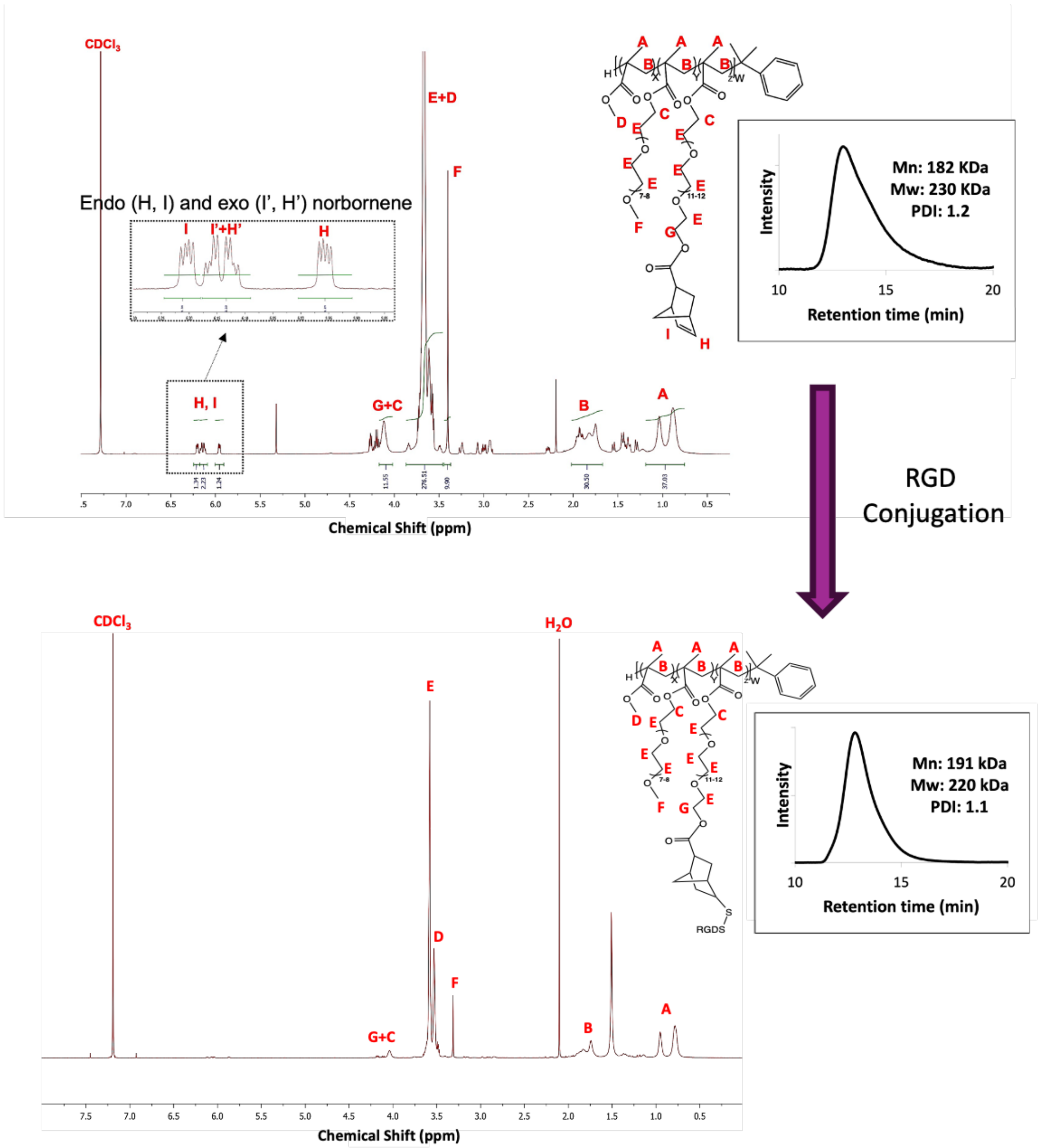


**Figure S2.** $^{1}$H NMR spectrum in deuterated chloroform and GPC spectrum in DMF from MPP polymer after esterification with norbornene and after thiol-ene click with peptide. The main NMR picks related to the conjugation of norbornene groups appear around 6.0 ppm, which is attributed to endo/exo protons that, after peptide conjugation, disappear.

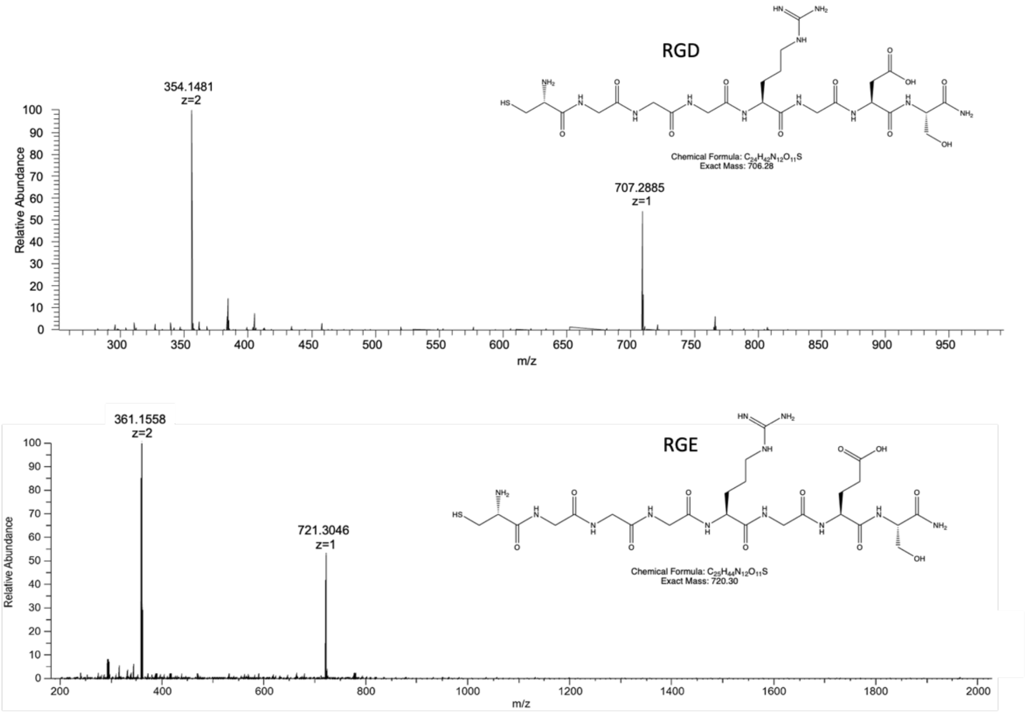


**Figure S3.** Mass spectrum of the purified RGD (Exact Mass: 706.28, Observed Mass: 707.28 (M+H), 354.14 (M+2H)), and RGE (Exact Mass: 720.30, Observed Mass: 721.30 (M+H), 361.15 (M+2H)).

After norbornene, conjugation $^1$H NMR ($CDCl_3$) results include δ 6.18-6.22 (dd, 3H), δ 6.10-6.17 (m, 3H), δ 5.90-5.97 (dd, 3H), δ 4.02 (s, 3H), δ 3.52-3.75 (m, 3H), δ 3.41-3.52 (m, 1H), δ 3.3 (s, 1H), δ 1.62-1.88 (m, 1H), δ 0.65-1.2 (d, 1H).

**Table S1** shows the composition of the NB functionalized polymers, as well as the formulae used to calculate the $^1$H NMR and GPC data. According to the formulae, $I_{PEGMA+PEGMANB}$ represents the total integral area of the PEG monomers inside the polymer. Furthermore, $I_{G+C}$ represents the integral area ascribed to the $CH_2$ peaks of the $COOCH_2$ methylene group protons in PEGMA+PEGMANB. The presence of 2/3 represents the ratio of $CH_{2,C}$ to the total number of $CH_{2,C}$ and $CH_{2,G}$ in PEGMA+PEGMANB. $I_{PEGMAOH}$ denotes the integral area connected to PEGMAOH, whereas $X_{PEGMAOH}$ and $X_{PEGMA+PEGMAOH}$ reflect the mole percentages of PEGMAOH and PEGMA+PEGMAOH in the polymer, respectively. These values are determined from the equations in **Table S1**. $X_{PEGMANB}$ represents the mole percentage of PEGMANB in the polymer. $I_H$ and $I_{I'+H'}$ denote the integral regions associated with norbornene's CH peaks in PEGMANB residues.

**Table S1.** Composition of polymers used in this study.

| Polymer name | Time of polymerization (h) | Conversion | Polymer composition before conjugation (mol%) (MMA-PEGMA-PEGMAOH) | Polymer composition after conjugation (mol%) (MMA-PEGMA-PEGMAOH-PEGMA-NB) | $M_n$ (kDa) | PDI |
|---|---|---|---|---|---|---|
| **MP** | 17 h | 60% | 84.4-15.6-0 | - | 128 | 1.1 |
| **MPP** | 17 h | 66% | 83.7-2.3-14.0 | 83.7-2.3-1.4-12.6 | 182 | 1.2 |

The actual polymer compositions were calculated based on H NMR data using the equations below:

**MP:**

$$X_{PEGMA} = \frac{\frac{I_C}{2}}{\frac{I_A}{3}} \times 100 \qquad X_{MMA} = 100 - X_{PEGMA}$$

**MPP:**

$$I_{PEGMA+PEGMANB} = \frac{I_{G+C}}{2} \times \frac{2}{3}$$

$$X_{PEGMANB} = X_{PEGMAOH} \times \frac{I_H + \frac{I_{I'+H'}}{2}}{I_{PEGMAOH}}$$

$$I_{PEGMAOH} = I_{PEGMA+PEGMANB} \times \frac{X_{PEGMAOH}}{X_{PEGMA+PEGMAOH}}$$

It is challenging to quantify peptides inside functional polymers using NMR because the associated peaks fall below the detection threshold of the NMR spectra. As a result, the peptide density for CHNS was estimated using trace elements microanalysis (conducted by Macquarie University Analytical and Fabrication Facility, School of Natural Sciences, NSW, Australia). In this application, bulk peptide density refers to the density of peptide in the bulk (measured in µg per mg of polymer). **Equation 1** was used to do the computation. Here, Y represents the nitrogen concentration of the polymer (measured in mg of nitrogen per kg of polymer), as determined by trace elements microanalysis. $M_{peptide}$ is the molecular weight of the peptide, $M_N$ is the molecular weight of nitrogen, n is the number of nitrogen atoms per mole of peptide (n = 12), and $10^3$ is included for unit conversion to µg/mg.

Furthermore, the number of peptides per polymer chain (Peptides/Chain) was calculated using equation (2). In this equation, $M_{nPolymer}$ represents the polymer's number-average molecular weight as determined by GPC analysis, and $10^6$ is the unit conversion used to calculate the number of peptides per polymer chain. The results are shown in **Table 1**.

$$Bulk\ peptide\ density = \frac{Y \times M_{Peptide}}{M_N \times n \times 10^3} \quad (1)$$

$$Peptides/Chain = \frac{Y \times M_{n\ Polymer}}{M_N \times n \times 10^6} \quad (2)$$

### 2.3. Optimization of electrospinning parameters for the polymer solution

In order to prepare nanofibrous samples in both aligned and random orientations, optimization of spinning parameters has been done. Accordingly, solutions of polymer in a mixture of methanol-water with different concentrations of 3, 6, and 10 wt.% were prepared. Then, the solutions were electrospun using a blunt 18G needle on a static stainless steel plate with different spinning voltages (12 and 17 kV) and feeding rates (0.6 and 1 $ml.h^{-1}$) while keeping the distance between the needle and collector constant at 15 cm. The spinning parameters were optimized based on the data in **Table S2,** in which the condition 11 parameters resulted in uniform nanofibers. For random and aligned fibers, the rotating collector with a rotation speed of 100 and 800 rpm was used, respectively.

**Table S2.** Optimizing the electrospinning parameters for the preparation of nanofibrous samples.

| **Condition** | **Polymer concentration (wt.%)** | **Flow rate ($ml.h^{-1}$)** | **Voltage (kV)** | **Distance (cm)** | **Needle gauge** | **Result** |
|---|---|---|---|---|---|---|
| 1 | 3 | 0.6 | 12 | 15 | 18 | Particle |
| 2 | 3 | 1 | 12 | 15 | 18 | Particle |
| 3 | 3 | 0.6 | 17 | 15 | 18 | Particle |
| 4 | 3 | 1 | 17 | 15 | 18 | - |
| 5 | 6 | 0.6 | 12 | 15 | 18 | Particle |
| 6 | 6 | 1 | 12 | 15 | 18 | Mixed fibers and particle |
| 7 | 6 | 0.6 | 17 | 15 | 18 | Mixed fibers |

| | | | | | | |
|---|---|---|---|---|---|---|
| | | | | | | and particles |
| 8 | 6 | 1 | 17 | 15 | 18 | Particle |
| 9 | 10 | 0.6 | 12 | 15 | 18 | Beaded fibers |
| 10 | 10 | 1 | 12 | 15 | 18 | Mixed fibers and particles |
| 11 | 10 | 0.6 | 17 | 15 | 18 | Uniform fiber |
| 12 | 10 | 1 | 17 | 15 | 18 | Fiber |

#### 2.4. NHS-NPs nanoparticles functionalization of surfaces

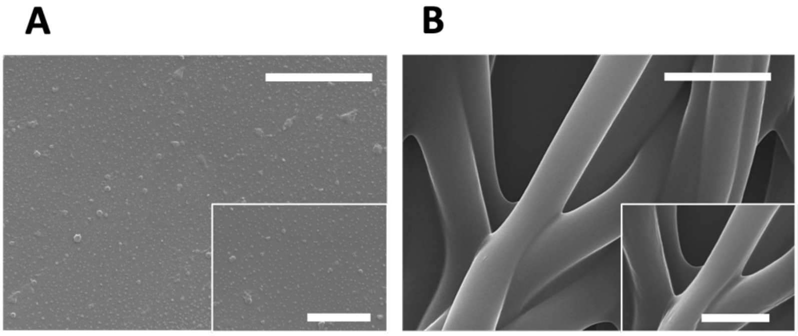


**Figure S4.** SEM images of A) NHS-AuNPs and B) L0G0 fiber surface after NP conjugation (scale bars are 3 and 1 µm for big and small panels, respectively.

#### 2.5. Evaluation of the morphology of the random fibers after 7 days of incubation

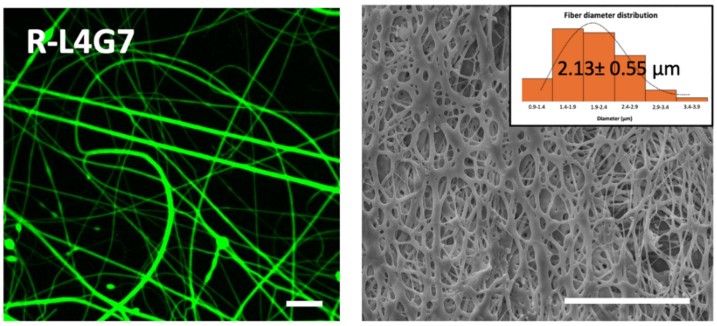


**Figure S5.** Representative images of random fibers (R-L4G7) after 7 days of incubation in PBS at 37°C, including (left) a confocal microscopy image of FITC-incorporated fibers (scale bar: 50 µm) and (right) an SEM image with fiber diameter distribution histograms (scale bar: 50 µm).

### 2.6. Evaluation of cell adhesion on different morphologies and ligand density

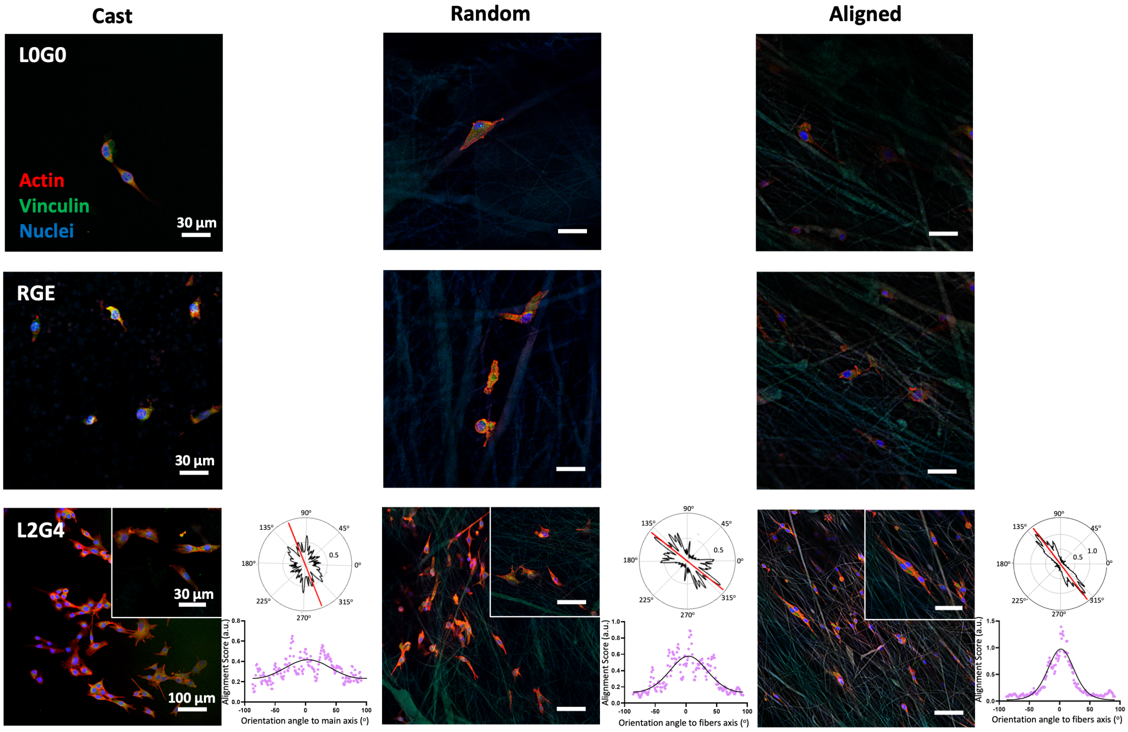


**Figure S6.** Representative confocal microscopy images of cells cultured on L0G0, RGE, and L2G4 surfaces after 24 h with cellular alignment analysis. Staining was done for actin filaments (red), vinculin (green), and nuclei (blue).

### 2.7. Evaluation of myogenesis and myotube maturation after 7 days of differentiation

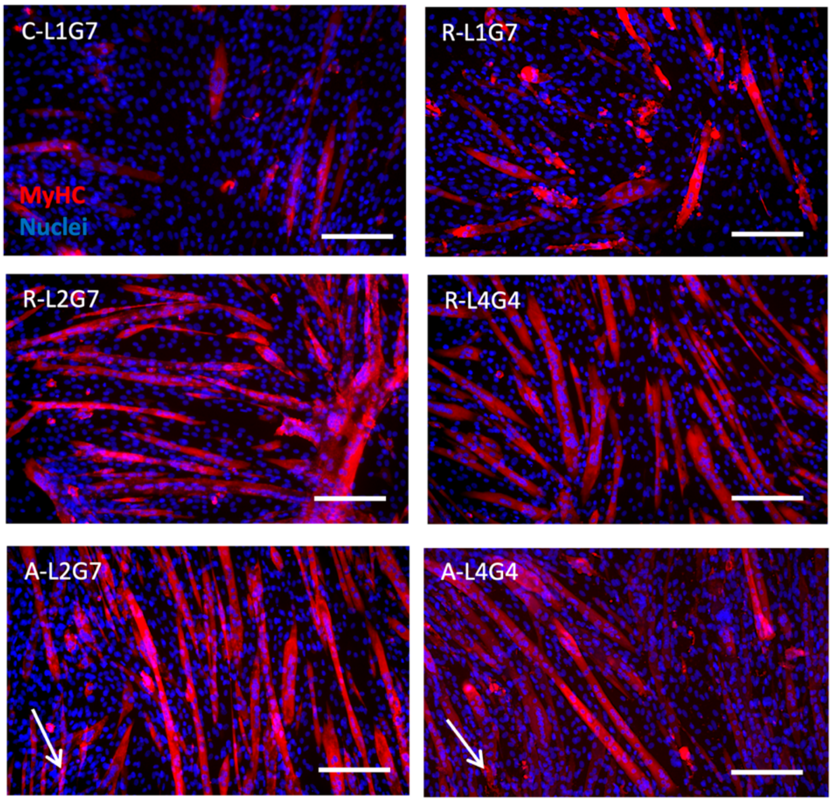


**Figure S7.** Assessment of myogenesis after 7 days of differentiation on various biomaterial samples. Representative confocal microscopy images of myotubes against MyHC (red) and nuclei (blue) (Scale bar is 200 µm and white arrows show the direction of fiber alignment).

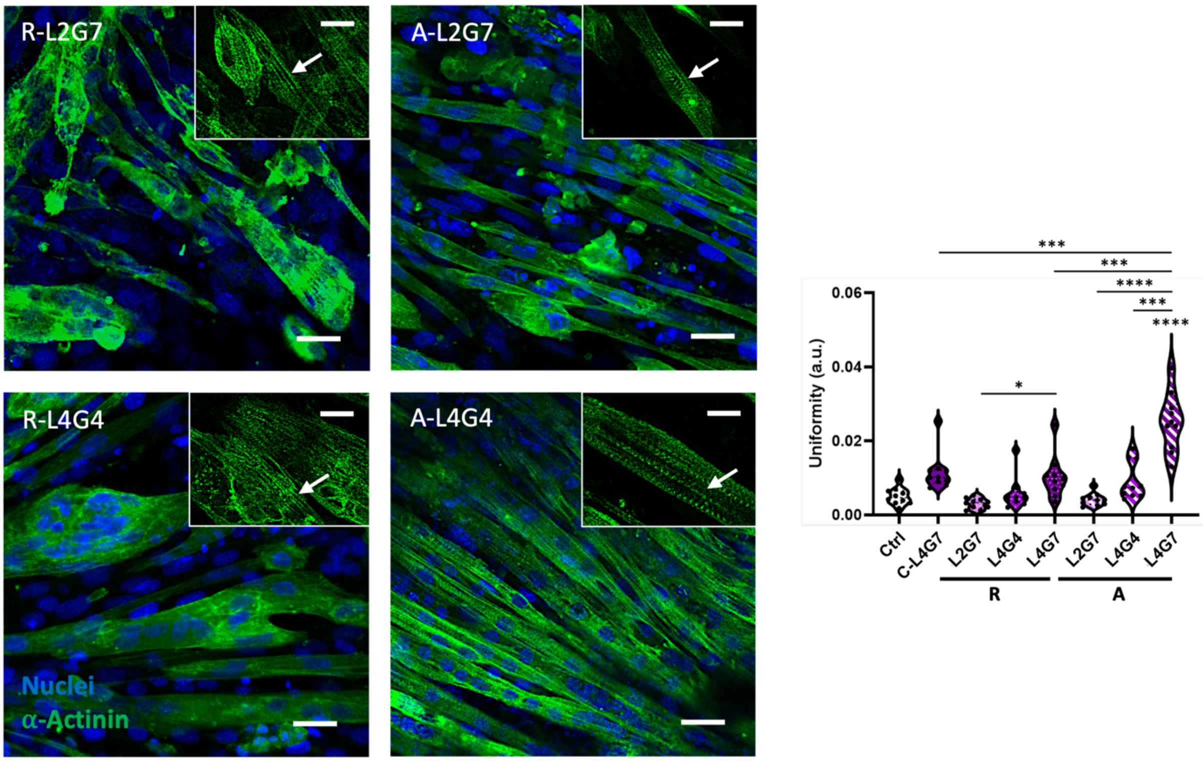

**Figure S8.** Assessment of maturation after 7 days of differentiation on various biomaterial samples. Representative confocal microscopy images against sarcomeric $\alpha$-actinin (green) and nuclei (blue) reveal the formation of z-lines and striation of myofibers, indicating their maturation (indicated by white arrows, scale bar of 50 µm). Images were further assessed to measure sarcomere uniformity using SotaTool software (n=10). Asterisks (*) directly above data points indicate statistical differences with the control, and asterisks above the bars indicate statistical differences between treatment groups.

#### 2.8. Evaluation of motor neuron differentiation

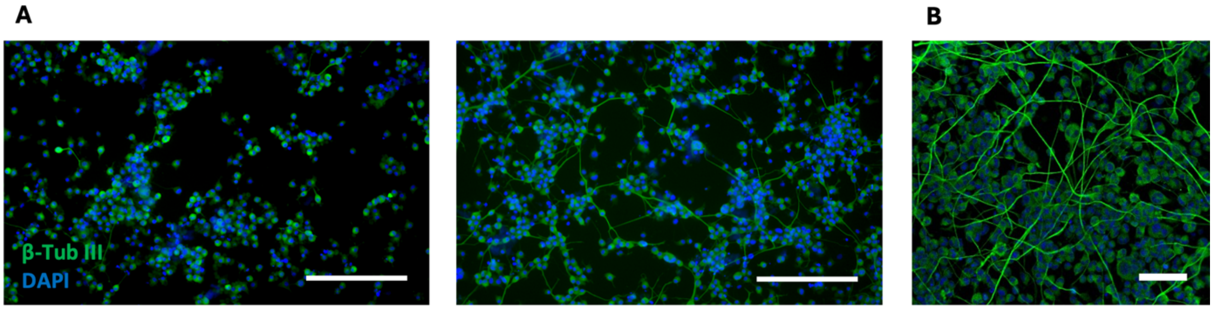


**Figure S9.** Representative confocal image of motor neurons A) before (left) and after (right) differentiation, before co-culture, and B) representative confocal image of motor neurons after differentiation on fibronectin-coated coverslip as positive control (staining against β-tubulin III (green), and nucleus (blue), scale bars for A and B are 200 and 50 µm, respectively).

#### 2.9. Analysis of contraction and NMJ functionality

**Table S3.** Description of the supporting information movies corresponds to the graphs in **Figure 9**.

| Movie code | Cell type | Substrate information | Added chemical | Results |
|---|---|---|---|---|
| **SIM-1** | C2C12 myoblasts | Control | - | Limited spot contraction |
| **SIM-2** | C2C12 myoblasts | A-L4G7 | - | Spontaneous contraction |
| **SIM-3** | C2C12 myoblasts co-cultured with NSC-34 motor neurons | A-L4G7 | - | Uniform contraction |
| **SIM-4** | C2C12 myoblasts co-cultured with | A-L4G7 | 10 µM acetylcholine | Triggered contraction (semi-tetanus) |

| | | | | |
|---|---|---|---|---|
| | NSC-34 motor neurons | | | |
| **SIM-5** | C2C12 myoblasts co-cultured with NSC-34 motor neurons | A-L4G7 | 10 μM tubocurarine | Inhibited contraction |
